# Rapid Detection and Initial Treatment of Life-Threatening Subdural Hematoma in a Sheep Model

**DOI:** 10.64898/2026.11.03.763711

**Authors:** David L Brody, Bradley A. Dengler, Joseph P. Angelo, David W. Blodgett, Adam B. Cohen, David G. Drewry, Michael J. Fitch, Nathaniel Goonasekeram, Adolph J. Januszkiewicz, Taylor Knapp, Brian Kohler, Joe Mark, Ian S. McAdams, Kathleen M. Perrino, Carissa L. Rodriguez, Timothy A. Sleasman, Graham L. Spicer, Nicole E. Steiner, Austin P. Veith, Jeremiah J. Wathen, Kevin C. Wolfe

## Abstract

Traumatic brain injury (TBI) is the leading cause of death and disability in people under 40 worldwide. There is an unmet need for detection and treatment of life-threatening TBI when computed tomography (CT) scanners and neurosurgeons are unavailable. We developed a new handheld frequency-domain multidetector infrared hemorrhage detection device. Using the new device, military pre-hospital medics required <4 minutes to detect life-threatening subdural hematoma in a sheep model. We developed a second device that allows non-neurosurgeons to safely create a burr hole and partially remove clotted subdural blood. In sheep, the treatment device reduced intracranial pressure within 20 minutes, and improved post-anesthesia blinded neurological outcomes. With human use adaptation, these devices could allow rapid detection and initial treatment of some life-threatening TBIs in challenging situations.

## Main Text

There are an estimated 21 to 69 million people who sustain a traumatic brain injury (TBI) every year (*1, 2*). TBI ranges in severity from concussion to life-threatening or fatal injuries (*3*). TBI is the leading cause of death and disability in people under 40 worldwide (*1, 4, 5*). On the more severe end of the spectrum, up to half may have bleeding underneath the skull but outside of the brain itself (*6*). Bleeding underneath the dura, the leathery coating of the brain, defines subdural hematoma. Bleeding outside the dura but under the skull defines epidural hematoma. Subdural hematoma is more common than epidural hematoma, but both types of hematomas can dramatically increase intracranial pressure (ICP). Increased ICP can cause reduced blood flow to the brain and other complications, which can result in permanent brain damage, persistent vegetative state and death (*7, 8*). The standard of care for severe TBI with subdural or epidural hematoma involves rapid detection and localization using a CT scan, followed often by immediate brain surgery to remove the clotted blood and reduce ICP (*9,10*). Immediate treatment can greatly improve outcomes for subdural and epidural hematomas (*9,11*). However, there are many situations where CT scanners and neurosurgeons are not immediately available (*12,13*). In large scale combat operations such as those ongoing in Ukraine (*14–16*), mass casualties, natural disasters, or in peer or near-peer conflicts, it may not be possible to rapidly transport all injured personnel to appropriate medical facilities (*17*). Furthermore, rural, wilderness, and ship-board areas can be many hours away from medical care even by air transport (*18–20*). In much of the developing world, adequate medical facilities may not be available (*21–23*) and mortality may be elevated 3-4-fold (*24*).

Several technologies have been developed to identify serious TBI without the use of CT scanning. These include blood biomarkers of brain injury, automated eye movement tracking devices, quantitative electroencephalogram (EEG), and hand-held infrared hemorrhage detection devices. The blood biomarkers, eye movement and EEG devices are designed for triage, but do not provide sufficient information to guide invasive treatment; specifically, they do not differentiate surgically treatable subdural and epidural hematomas from other types of TBIs. The existing hand-held infrared hemorrhage detection devices can provide some information about the location of subdural and epidural hemorrhages, but they have limitations (*25–27*). They compare the relative amplitude of the detected infrared for the right side vs. left side at fixed positions but cannot make independent assessments of each brain region (*25, 26*). Thus, they are not sensitive to bilateral subdural hemorrhages, which occur in about 15% of cases (*28*). Furthermore, the existing devices cannot differentiate between subdural and epidural hemorrhages, which can be life threatening, vs. scalp hemorrhages, which are not life threatening. And finally, they do not localize hemorrhages precisely enough to guide treatment (*29, 30*). Thus, there is an unmet need for solutions to detect and localize subdural and epidural hemorrhage with sufficient resolution to guide treatment when CT scanning is not available.

## RESULTS

### Infrared Intracranial Hemorrhage Detection Device

With guidance from US Special Forces medical personnel, we developed a new frequency domain multi-detector infrared intracranial hemorrhage detection device **(Fig. 1** and **Fig. S2)**. The new device incorporated 3 advanced methods to allow independent assessment of each brain region with higher spatial resolution and differentiate intracranial (e.g. subdural and epidural) blood from superficial scalp hemorrhage. First, the new device employed 3 detectors instead of one. The detectors more distant from the source are expected in principle to be more sensitive to infrared light scattered from deeper tissues whereas the closer detectors are expected to be more sensitive to light scattered from more superficial tissues **(Fig. 1A)**. Second, the new device included an infrared laser whose intensity was modulated rather than remaining constant. The modulated laser intensity allows information about the phase shift of the infrared light scattered by the tissues between the laser source and the detectors to be used in addition to the amplitude **(Fig. 1B)**. Third, the new device was designed to be pressed against the head and moved around continuously, rather than applied only at fixed positions. This provided more information about the location and extent of the intracranial hemorrhage ( **Fig. 1C)**. The position of the device was tracked and signal quality was monitored to ensure adequate coverage of the accessible parts of the intracranial space **(Fig. 1D)**. The new device was tested and calibrated using optical phantoms with characteristics similar to those of human scalp, skull, and blood **(Supplemental Methods)**.

**Fig. 1:**
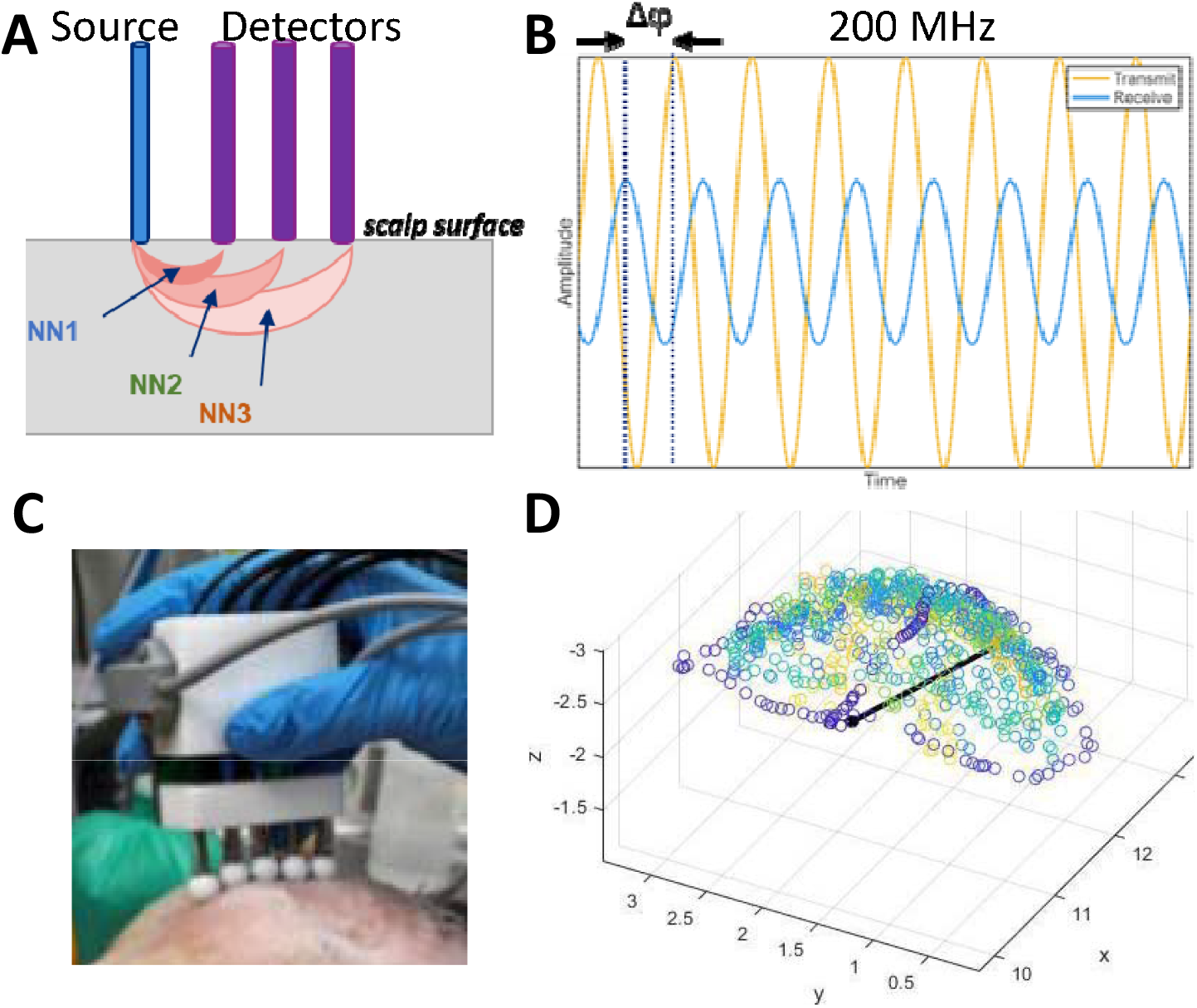
Handheld frequency domain infrared intracranial hemorrhage detection device. **(A)** Conceptual design depicting the geometry of the transmit and receive channels on handheld probe used for measurement, with detector fibers spaced spaced 1, 2, and 3 cm from the source. The light scattered from more superficial tissues is differentially detected at nearest neighbor 1 (NN1) whereas light scattered from deeper tissues is differentially detected at nearest neighbors 2 and 3 (NN2, NN3). (**B**) Amplitude of the transmitted infrared light is modulated at 200 MHz. Both phase shift (Δϕ) and amplitude attenuation are measured at each detector. (**C**) Prototype handheld detection device involving a 850 nm laser infrared light source and silicon photomultiplier detectors tested on the shaved scalp of a sheep. (**D**) Spatial tracking of the handheld detection device over the surface of the head, with data quality feedback to the user. Blue points indicate the outline of the coverage area, and other points indicate coverage density within the coverage area.

### Sheep Model of Life-threatening Subdural Hematoma

To test the new detection device *in vivo, we* developed a sheep model of life-threatening subdural hematoma based on previous pig models (*31, 32*) under an Institutional Animal Care and Use Committee (IACUC) approved protocol **(Fig. 2)**. Sheep were chosen because they have been used in TBI research successfully (*33, 34*) and their skull thickness and optical properties are similar to those of humans **(Fig S1)**. Pig skulls are typically too thick, and testing involving non-human primate species was not considered ethically acceptable. Subdural hematoma was created by instilling fresh autologous venous blood into the subdural space under propofol anesthesia in steps of 1 mL/minute **(Fig. 2A)**. The subdural space was accessed through a small catheter placed through a groove drilled in the skull overlying the left frontal cortex **(Fig. S3)**. CT scans of the head **(Fig. 2B** and **Fig. S4-S5)** and post-mortem examinations **(Fig. 2C** and **Fig. S6)** were used to verify the location of the subdural hematoma. To calibrate the severity of the model, we tested infusions of 10,12,15, and 20 mL of blood. Our goal was to create a model with 50% mortality, similar to the natural history of large subdural hematoma in human patients with severe TBI. Mortality after 15-mL infusions was 3/6, whereas it was lower with 10- and 12mL infusions and higher with 20-mL infusions **(Fig. 2D)**. Therefore, 15-mL infusions were used for subsequent subdural hematomas. Deaths were mainly due to apnea or severe neurological deficits requiring humane euthanasia. Neurological examinations using a 24-point scale **(Fig. S7)** indicated a range of impairments at 72 hours, with severity increasing for sheep with infusions of larger volumes of blood **(Table S1)**. ICP was measured using an intraparenchymal pressure catheter placed contralaterally to the subdural catheter **(Fig. 2A)**. ICP was elevated in a nonlinear fashion; early blood infusions caused modest increases in ICP, reflecting maintained compliance, while later blood infusions caused more dramatic increases in ICP, indicating reduced compliance **(Fig. 2E)**. The mean baseline ICP was 12 mm Hg and mean peak ICP was 114 mm Hg, falling to a mean of 85 mm Hg after 5 minutes **(Fig. 2F)**. Detailed ICP **(Fig. S8)**, hemodynamic **(Fig S9-10)**, and respiratory **(Fig. S11** and **Table S2)** measurements indicated physiological responses like those seen in humans with TBI. Taken together, these findings indicated that the sheep subdural hematoma model had properties similar to life-threatening subdural hematoma in human patients with TBI. This model may be useful for many other pathophysiological, diagnostic, and therapeutic investigations **(Supplementary Discussion)**.

**Fig. 2:**
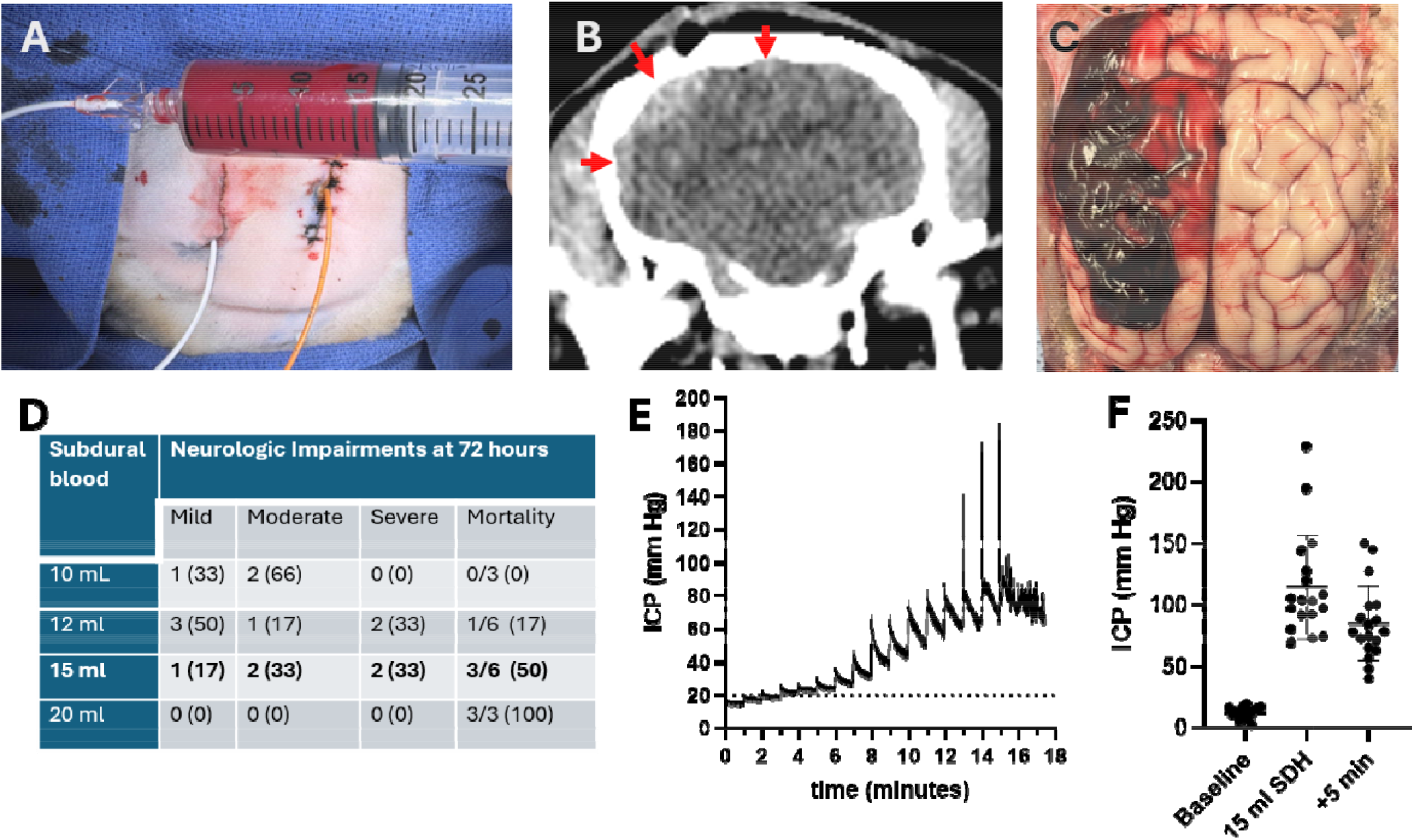
Sheep model of life-threatening subdural hematoma. **(A)** Infusion of autologous venous blood through a subdural catheter (white) with a parenchymal fiber optic ICP monitor implanted contralaterally (yellow). (**B**) CT scan showing subdural hematoma, indicated by red arrows. (**C**) Post-mortem photograph of sheep brain after removal of scalp and dura, showing subdural hematoma. (**D**) Neurological impairments after 10-, 12-, 15-, and 20-mL infusions of subdural blood. Mild: rapid recovery from anesthesia and surgery with minimal observable effects and surviving 72 hours. Moderate: Recovery from anesthesia and surgery residual neurological deficits but surviving 72 hours. Severe: Debilitating effects lasting ≥24 hours, incompatible with normal life functioning. Animals in this group were generally subject to early euthanasia. The one exception was one sheep in the 12-mL group with severe neurological deficit but ultimately survived the 72-hour observation period. Criterion of 50% mortality was met for 15-mL blood infusion. Neurological outcomes could not be measured at 72 hours for all sheep because of early mortality. (**E**) ICP as a function of time during bolus infusions of 1 mL of blood each minute for 15 minutes through the subdural catheter. (**F**) ICP at baseline, immediately after infusion of 15 mL of blood, and 5 minutes later (n=18).

### Accuracy of Intracranial Hemorrhage Detection Device

Next, we tested the accuracy and usability of the new intracranial hemorrhage detection device in the sheep model **(Fig. 3)**. An expert user calibrated the device and performed scans of each sheep at baseline before any intracranial interventions were performed to verify that there were no anatomical abnormalities that could distort the subsequent results **(Fig. 3A)**. Then, 23 sheep were randomly assigned to either subdural hematoma or sham. The sham procedures involved scalp incisions but no skull groove or subdural blood infusion. All catheters were removed and skin was sutured identically **(Fig. S3G)**. We trained 8 medical providers with skills and training comparable to intended real-world users (5 Army medics, 1 Navy corpsmen, 1 SEAL medic, 1 developing world Emergency Medicine physician). Training involved 30 minutes of theory and hands-on practice with the device. Typically, 2 medical providers assessed each sheep with the new device in a blinded fashion. The medical providers were able to obtain good quality scans in 3-4 minutes **(Fig. 3B)**. Sensitivity for subdural hematoma using the new infrared hemorrhage detection device was 0.58 and specificity was 0.73, with an overall accuracy of 0.67. A different set of medical providers used the currently approved Infrascanner 2000 device to assess the same sheep for 3-4 minutes in a blinded fashion. Sensitivity was 0.45 and specificity was 0.84, with an overall accuracy of 0.67. Thus, for a relatively simple scenario, unilateral subdural hemorrhage vs. no hemorrhage, the new intracranial hemorrhage detection device had similar overall accuracy as the currently approved Infrascanner 2000 device (*29, 30*) in a sheep model.

**Fig. 3:**
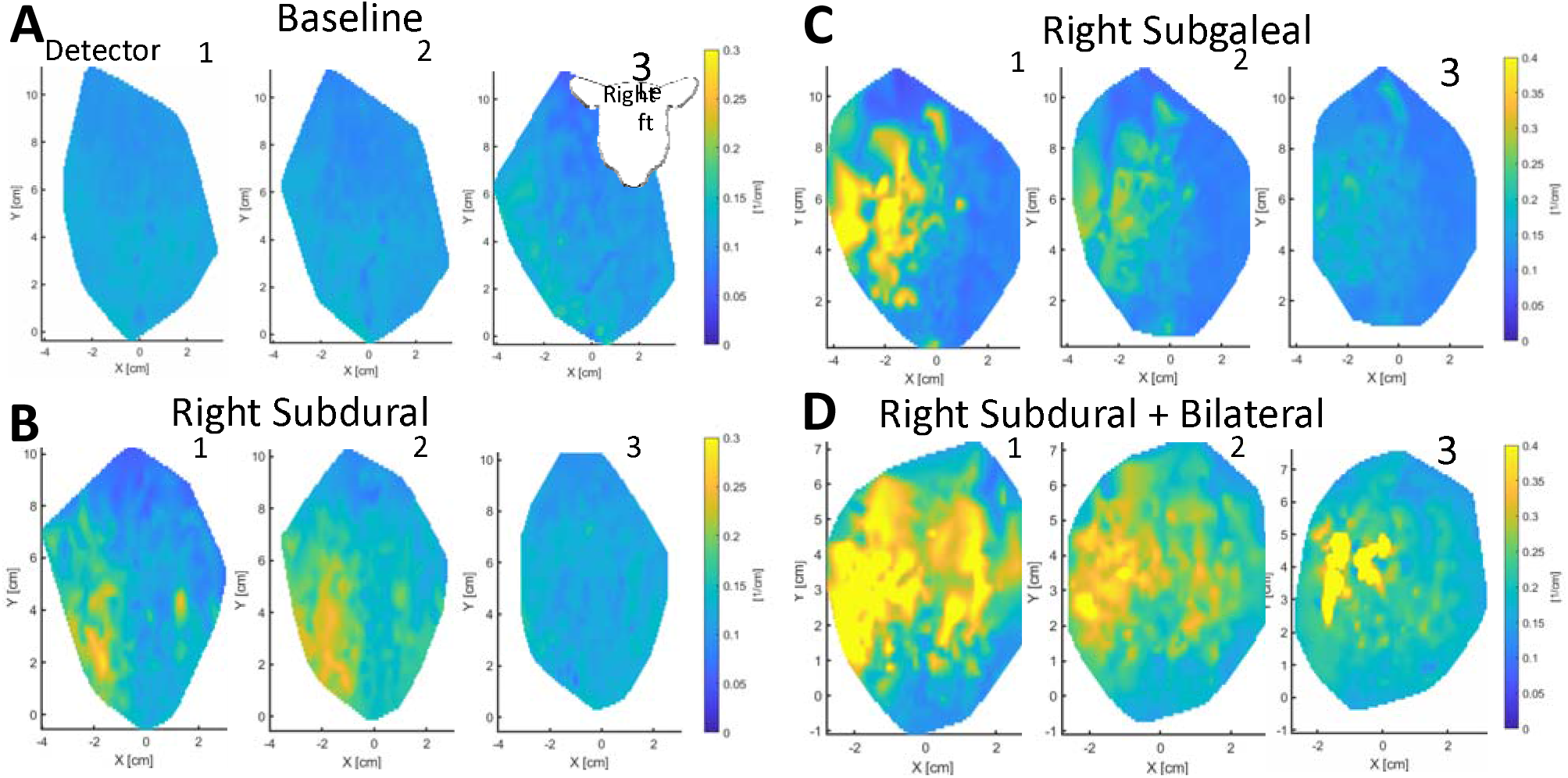
Infrared detection of subdural hematoma in a sheep model. Spatial maps of infrared amplitude attenuation for detectors 1, 2, and 3, with warmer colors indicating greater attenuation (**A**) Baseline before any intracranial procedures showing relatively low infrared attenuation throughout. (**B**) Right-sided subdural hematoma showing infrared attenuation on the left in all 3 detectors. (**C**) Right-sided non-life-threatening subgaleal blood showing extensive attenuation in detectors 1 and 2, but less attenuation in detector 3. (**D**) Combination of Right - sided subdural hematoma and bilateral subgaleal blood showing extensive bilateral attenuation in detectors 1 and 2, with greater attenuation in detector 3 on the right (with subdural and subgaleal) than on the left (with subgaleal only).

However, under more complex scenarios, the medical providers were more accurate using the new intracranial hemorrhage detector than using the Infrascanner 2000. As noted above, superficial scalp hemorrhages commonly occur after head trauma. While not life threatening, blood in the scalp attenuates infrared light and may complicate detection of subdural hematoma both by creating false positives when no subdural exists and obscuring subdural hemorrhages when they are present. Some superficial scalp hemorrhages are readily visible by eye and can be avoided, but others in the deeper subgaleal layer may not be readily visible. Therefore, after testing for subdural hematomas as described above, we subsequently randomized the same sheep to receive ipsilateral, contralateral, bilateral or no subgaleal autologous blood infusions **(Fig. S6H)**. We used the same sheep to reduce the total number of animals used. For subgaleal blood infusions, 2 mL of autologous blood was injected in a fan pattern using a 25-gauge 1 inch needle. Shams had 25-gauge needle sticks but no blood infusion and were visually indistinguishable *in vivo*. The subgaleal blood attenuated the infrared light received by the first detector of the new intracranial hemorrhage detection device **(Fig. 3C)**, but there was less attenuation in the more distant detectors, as expected based on the underlying principles **(Fig. 1A)**. When both subdural and subgaleal hemorrhage were present, a complex mixture of signals was observed **(Fig. 3D)**. In these complex cases involving possible subgaleal blood, subdural hemorrhage, both, or neither, the medical providers had a sensitivity of 0.84, specificity of 0.56, and unchanged overall accuracy of 0.66 for detection of lifethreatening subdural hematoma using the new infrared hemorrhage detection device. In contrast, sensitivity was 0.38, specificity was 0.65, and overall accuracy was 0.55 with the Infrascanner 2000. For the new infrared hemorrhage detection device, use of multiple modulation frequencies from 100 MHz to 500 MHz and use of a support vector machine learning classifier provided comparable results **(Supplemental Results and Table S3)**. For bilateral subdural hematomas, the new infrared hemorrhage detection device had an overall accuracy of 0.87, whereas the Infrascanner 2000 overall accuracy was 0.38. Thus, the new infrared hemorrhage detection device was able to address several of the limitations of the Infrascanner, including localization of the extent of subdural hematoma, detection of subdural hematoma in the presence of subgaleal blood, and accurate detection of bilateral subdural hematomas.

### Initial Treatment of Subdural Hematoma

Detection of life-threatening subdural hematoma would be most useful when paired with rapid, safe, and effective initial treatment. At present, non-neurosurgeons generally lack the capabilities to provide such treatment (*35, 36*). Anecdotal evidence indicates that in wartime (B. Moore, R. Thomas, and J. Holcomb, personal communications), other remote settings, or acute emergencies (*35, 37, 38*) some non-neurosurgeon health care providers have drilled emergency burr holes through the skull to attempt to remove some subdural and epidural blood. It is not clear how safe and effective these treatments have been (*36, 39*). Importantly, intracranial blood typically clots rapidly, and it can be difficult to remove clotted blood through a relatively small burr hole. Furthermore, in a non-sterile environment, creation of an open burr hole puts the patient at elevated risk of infections such as meningitis and encephalitis, both of which can also cause permanent neurological damage and death. Thus, standard open burr hole treatments are not considered a substitute for definitive neurosurgical treatment and currently are not part of standard care even in challenging circumstances (*40*).

Again, with consultation from US Special Forces medical personnel and neurosurgical colleagues, we designed a second hand-held device that allows non-neurosurgeons to safely create a burr hole and remove some of the clotted subdural blood using a morcellator **(Fig. 4)**. The device included 5 main features **(Fig. 4A** and **Fig. S12-S14)**. First, it was designed to attach securely to the head via 3 bone screws, pressing a foam rubber gasket tightly against the skin as a barrier to infection. For treatment of life-threatening TBI, the user would shave the head and clean the scalp thoroughly before fixing the device to the patient’s head overlying the subdural or epidural hematoma. Second, it included a battery-powered cautery ring to burn through the scalp and prevent superficial scalp bleeding. Third, it included an approximately 1 cm diameter hand crank drill with a pressure-sensitive clutch to limit the axial force applied and a hard stop to prevent it from advancing past the inner table of the skull. This feature was designed to reduce the risk of drilling into the brain itself. Fourth, it included a deployable blade to cut the dura. Fifth, it included a transparent lid with an access port through which a morcellator could be passed through the burr hole and into the intracranial space. It was designed to be compatible with a portable version of the hand-held NICO Myriad morcellator device which applies suction and an alternating cutting blade to remove clotted blood (*41, 42*).

**Fig. 4:**
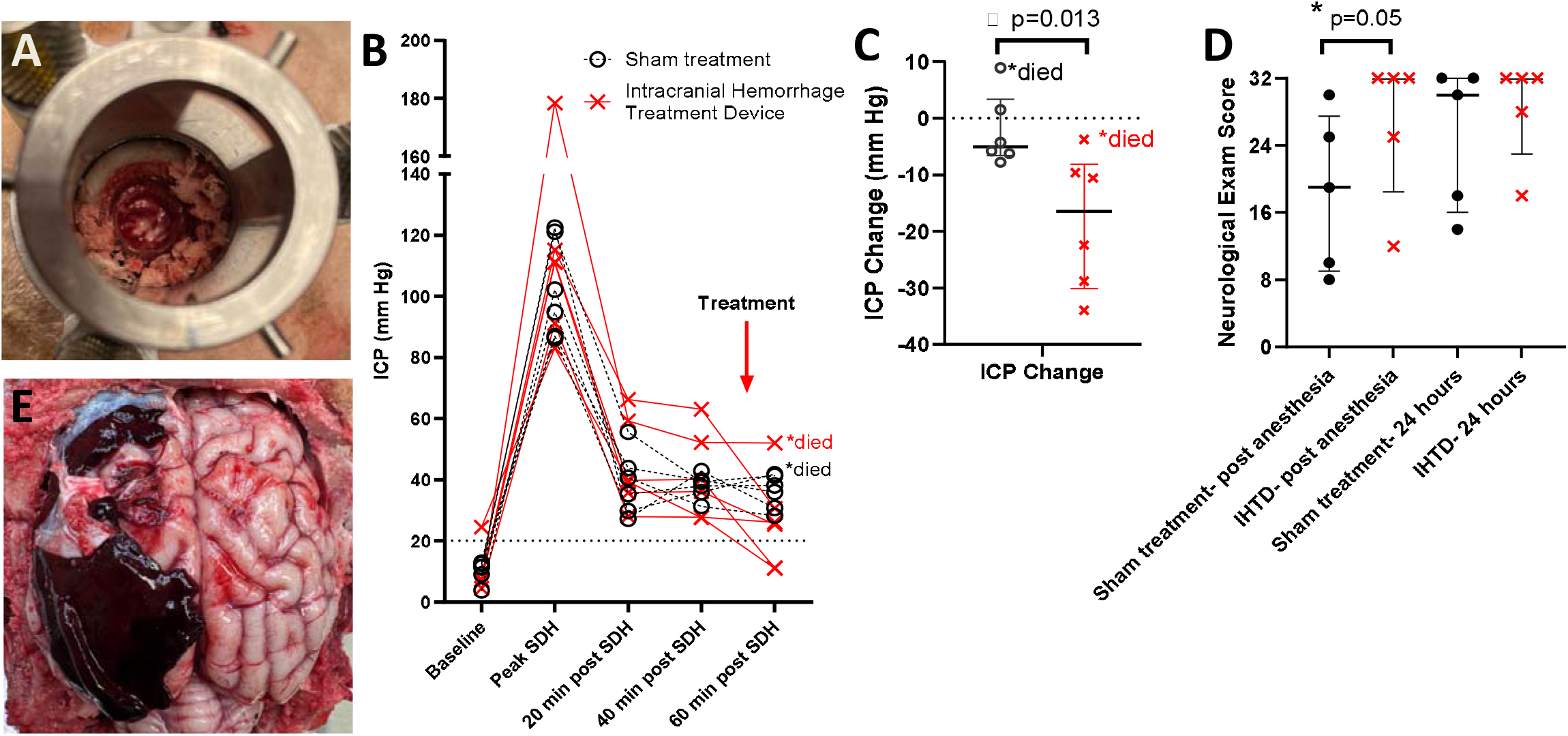
Treatment of life-threatening subdural hematoma in a sheep model. **(A)** Treatment device base anchored to the head of a sheep, with a completed 1 cm burr hole and dural puncture revealing underlying subdural hematoma. (**B**) ICP at prespecified time points: baseline, at peak following subdural hematoma creation, 20 to 40 minutes following subdural hematoma creation but before treatment or sham, and 60 minutes following subdural hematoma creation after treatment or sham. Each ICP measurement was made by averaging 1 minute of artifact free ICP recordings sampled at 100 Hz. (**C**) Change in ICP from the average of the 20- and 40minute time points to the post-treatment or sham time point. *indicates p=0.013, 1-sided Mann-Whitney U Test. (**D**) Neurological status assessed by blinded veterinarians after recovery from anesthesia approximately 3-5 hours after subdural hematoma creation, and again 24 hours after subdural hematoma creation. *indicates p=0.05, 1-sided Mann-Whitney U Test. (**E**) Postmortem photograph showing subdural hematoma anterior and posterior to a cleared region where the hematoma was removed using the morcellation device (compare with Fig. 2C and Fig. S6)

We performed a preregistered, randomized, sham-controlled trial in sheep with a prespecified sample size of 12 qualifying sheep (https://osf.io/yvkqf/overview). We started with a sample size of 18 sheep and planned to exclude approximately 1/3 of the sheep with non-subdural intracranial blood (e.g. intraparenchymal, intraventricular: **Fig. S5-S6)**. The prespecified primary outcome measure was reduction in ICP. The trial had statistical power of 0.8 to detect a reduction in ICP with Cohen’s d effect size of 1.5 or greater using a 1-tailed student t-test at p<0.05, and similar power for a 1-sided Mann-Whitney U test in case of non-normally distributed data. Sheep were randomized after the creation of the subdural hematoma and initial ICP measurements to reduce bias. In the trial, ICP elevations were similar to those seen in earlier studies, and ICP 20-40 minutes after completion of the subdural hematoma ICP did not differ between groups **(Fig. 4B)**. Treatment time required under 20 minutes for a nonneurosurgeon, including removing the device from its sterile packaging, affixing it to the sheep head, cauterizing the scalp, drilling the burr hole, cutting the dura, and morcellating the subdural clotted blood **(Fig. S16)**. Use of the treatment device starting approximately 40 minutes after creation of the subdural hematoma reduced intracranial pressure by a mean of 16.5 mm Hg, whereas ICP declined by a mean of 5 mm Hg in the sham group during the same time period **(Fig. 4C**, p=0.013,1-sided Mann-Whitney U Test, rank biserial correlation r=0.78, large effect size equivalent to Cohen’s d=2.48). There were two device performance failures involving the dural cutting blade, both of which were corrected on-site after which normal operation resumed. There were no device-related adverse events. After final ICP measurements were performed, the device was removed and the sheep’s head was bandaged. Propofol anesthesia was stopped and sheep were transported to a recovery room. There was 1 early death in each group, both from apnea. The sheep that died early had greater than typical elevated ICPs after subdural blood infusion. Once the survivors had recovered from anesthesia, they were examined by veterinarians blinded to treatment status using an updated 32-point standardized neurological assessment **(Fig. S15)**. The surviving sheep in the treatment group had improved post-anesthesia neurological outcomes compared with the sham group **(Fig. 4D**, p=0.05,1-sided Mann-Whitney U test). Elevated final ICP was associated with impaired postanesthesia neurological function, with the correlation largely driven by the group of sheep randomized to treatment **(Fig. S17)**. There was some neurological recovery by 24 hours in many sheep in both groups **(Fig. 4D)**. All sheep were then humanely and respectfully euthanized. At post-mortem, the remaining subdural hematoma and zone where hematoma had been removed were clearly visible **(Fig. 4E)**. Taken together, these results indicate that the new treatment device could be used by a non-neurosurgeon to rapidly and effectively reduce ICP and improve neurological outcomes in a sheep model of life-threatening subdural hematoma.

## DISCUSSION

In summary, we have designed and tested two new prototype hand-held devices intended for detection and initial treatment of life-threatening subdural hematoma in challenging clinical environments. The devices were tested in a novel sheep model of life-threatening subdural hematoma. These devices are not intended to substitute for CT scanners and neurosurgeons, but when CT scanners and neurosurgeons are not available, these hand-held devices may facilitate diagnosis and initial management of life-threatening subdural hematoma. While not explicitly tested, it is likely that detection and treatment of epidural hematoma would be similarly facilitated. The two devices offer flexibility in their application with multiple potential implementation approaches by using them independently or in combination. We envision both devices being used together by front line medical providers in austere situations such as military operations, wilderness and shipboard medicine, and in many parts of the developing world. Severe TBI is dramatically more common in low- and middle-income countries with very limited access to high-level trauma care (*24, 43, 44*). In situations requiring rapid triage such as mass casualty events, the detection device could be used to determine who has the most serious subdural or epidural hemorrhage and has the highest need for immediate neurosurgical treatment. In situations where there may be a CT scanner but no neurosurgeon such as a level 3 or level 4 trauma center in the United States, an emergency medicine physician or general surgeon could use the treatment device based on the results of the CT scan. There are 493 level 3 and 923 level 4 trauma centers in the US (*45*). Many of them are in rural areas more than 1 hour away from a higher-level trauma center, and mortality from severe trauma can be substantially higher in rural environments (*46, 47*).

Before these results can be translated to clinical practice, a considerable amount of work is still required, including replication, longer term neurological outcome studies, infection risk assessments, assessment of optimal detection and treatment timing, miniaturization, ruggedization, Good Manufacturing Practice production, regulatory clearances, medical provider training, and human clinical testing. Nonetheless, the results presented here provide initial optimism regarding solutions to the unmet need for detection and initial treatment of life-threatening subdural hematoma when CT scanners and neurosurgeons are not immediately available.

## Acknowledgments

We are grateful to the Department of Laboratory Animal Medicine at Uniformed Services University, including Michael Bencivenga, James Bowring, Kelsey Fiddes, Ethan Holicky, Christopher Reeves, Kwasi Sampong, Katelyn Thomas, and Dawn Torissi; the veterinary team at TD Morris Inc., including Lynn Raab and Eleanor Lindauer; our forward military medical providers including Adrian Arintok, Micah Barton, Adam Brien, Meheret Fantaye, Reggie Jaramillo, Jason Shen, Dorian Snowden, Steve Viola, and TJ Zane; our military medical consultants including Randy Bell, Brock Benedict, Mike Davis, Patrick Depenbrock, John Holcomb, Todd Rassmussin, Stacy Shakleford, Andrew van Sumeren, and Steve Viola; technical assistance from Jared Wormley and Max Basescu; and surgical consultants Randy Bell, Nicholas Boulis, Justin Caplan, and John Willie.

## Funding

US Special Operations Command: W81XWH-22-C-00 (DLB)

Defense Health Agency Technology Transfer for the Warfighter Program: HU00012120079 (DLB)

Center for Neuroscience and Regenerative Medicine at the Uniformed Services University of the Health Sciences. CNRM-92-10010 (DLB)

## Author contributions

Conceptualization: DLB

Methodology: DLB, JPA, DWB, BAD, MJF, AJJ, JM, ISM, KMP, CLR, TAS, GLS, APV, JJW, KCW

Investigation: DLB, JPA, BAD, AJJ, BK, KMP, CLR, JJW, KCW

Visualization: DLB, AJJ

Funding acquisition: DLB

Project administration: DLB, DWB, ABC, DGD, TK, CLR

Supervision: DLB, BAD, AJJ

Writing-original draft: DLB

Writing - review & editing: DLB, AJJ, CLR

## Competing interests

A US Patent, US 12,150,655B2, David L. Brody, David Blodgett, Adam Cohen, Max R. Basescu, Kevin C. Wolfe, Jared M. Wormley. Date: Nov. 26, 2024, entitled “AUTOMATED CRANIAL BURR HOLE DEVICE AND METHOD” has been granted.

DLB serves as Editor-in-Chief *of Journal of Neurotrauma*

No other competing interests.

## Data, code, and materials availability

All raw data, code, and materials used in the analysis may be requested from the corresponding author. Device prototype availability will require materials transfer agreements (MTAs). All analyzed data are available in the main text or the supplementary materials.

## Disclaimers

The opinions and assertions expressed herein are those of the author(s) and do not reflect the official policy or position of the Uniformed Services University of the Health Sciences or the Department of War.

The views expressed herein are those of the author(s) and do not reflect the official policy or position of the Henry M. Jackson Foundation for the Advancement of Military Medicine, Inc. (HJF).

## Supplementary Materials

Supplementary Introduction

Materials and Methods

Supplementary Results

Supplementary Discussion Figs. SI to S17

Tables SI to S3

References (*1-56*)

