## SUPPLEMENT for "Rapid Detection and Initial Treatment of Life-Threatening Subdural Hematoma in a Sheep Model"

### SUPPLEMENTARY INTRODUCTION

Severe traumatic brain injury (TBI) is a leading cause of death and disability worldwide (*1*). In low- and middle-income countries, the incidence of severe TBI has risen with the use of motorized transport, while shifting demographics have resulted in more fall-related injuries in the aging population (*2*). In military personnel, exposure to blast is associated with unique pathological changes frequently accompanied by blunt force or penetrating trauma (*3, 4*). In the event of armed conflict with an advanced peer or near-peer adversary, severe injuries of all types are expected to be more common

than in recent wars and casualty care complicated by delayed medical evacuation (*5*). Although severe

injuries currently represent a small percentage of all TBI, they are associated with the highest levels of

morbidity and mortality. Among survivors, there is a reduced life expectancy compared to the general

population (*6*).

Severe TBI occurring in austere, rural, or remote settings presents special challenges above and

beyond those of the injury itself. Those living in rural areas have a higher annualized incidence of TBI,

and both total and prehospital mortality are higher when the injury occurs in a rural setting (*7, 8*). The fact that many rural hospitals in the U.S. are downsizing or closing exacerbates this problem. The number of Americans who live more than 1 hour away from trauma care has grown to 60 million (*9, 10*). This gap is of particular relevance to severe TBI for which rapid diagnostics and treatments are required. Level I trauma facilities are even more dispersed, and sub-specialists such as neurosurgeons are concentrated in urban areas. Areas without ready access to high acuity intensive care have been referred to as “trauma deserts” (*11*). In the military, expectations that future operational environments will require far-forward troops to delay medical evacuation and treat injured personnel onsite for extended periods has led to the concept of prolonged field care. Prolonged field care has become a major focus of US Military medical research and training in recent years (*12, 13*).

Intracranial extra-axial bleeding, including subdural hemorrhage (SDH) is associated with increased mortality in the setting of severe TBI (*14, 15*). Acute SDH complicates up to 11% of mild and moderate TBI and 50% of severe TBI (*14, 15*). Coma is present immediately in 25% to 50% of SDH patients, with another 12% to 38% progressing to coma in the following hours (*14*). Epidural hematoma, another form of intracranial extra-axial bleeding is less common and has generally more favorable prognosis. SDH is associated with worse prognosis both in developed nations (*16*) and after burr hole surgery in a more austere setting (*17*). The pathophysiology of SDH is characterized by widespread ischemia (*18*) and impaired cerebral blood flow, secondary to increased intracranial pressure (ICP) (*19*). This increased ICP is due in large part to the effects of the extra-axial mass lesions. Since the skull has a fixed volume and only enough space for blood, cerebrospinal fluid, and brain matter, any extra-axial blood will cause the ICP to rise dramatically after all compensation mechanisms have been exhausted. This can then lead to secondary brain injury through mechanisms of excitotoxicity (*20*) and apoptosis (*21*), and eventually herniation and death. Mortality associated with SDH varies depending on age, neurological status at the time of assessment, and injury severity (*22*) and in severe TBI, SDH is associated with greatly increased mortality rates, especially when caused by high velocity impacts (*23*). Even with high quality care as performed in the context of a clinical trial, overall mortality is greater than 30% (*24*). When treatment is delayed, mortality may reach as high as 80 – 90% (*25, 26*). Thus, patients with SDH can have poor outcomes and require urgent treatment.

Early surgical intervention may greatly reduce mortality and morbidity in patients with traumatic SDH (*26-29*). Many patients will require neurosurgical intervention to remove the mass lesion or decompress the brain, but in a military context these services are frequently limited and only available at Role 3 medical care. (Role 3 is defined as a fully staffed field hospital or equivalent). Additionally, monitoring and management of intracranial pressure can help drive aggressive medical interventions to stabilize patients awaiting transport to the Role 3 with Neurosurgical Capability (*30*). Thus, rapid identification and management is critical to reducing morbidity and mortality associated with SDH.

Most SDH is detected using CT scanning (*31*), but when CT scanners are not available, alternative methods are required. Near infrared spectroscopy (NIRS) and related techniques are well suited for noninvasive cerebral measurements. Near infrared light is able to penetrate through the scalp and skull to probe cortical tissue and provide insight into blood content and oxygenation by making estimates of the optical scatter and absorption properties of the tissue. In fact, handheld devices leveraging properties of NIR light propagation through biological tissue have been developed for intracranial hematoma detection (*32, 33*). These devices work by injecting NIR light into the tissue and detecting the amount of optical power that propagates to the detector. A comparison is made between the left and right side of the cranium at four different locations (frontal, temporal, parietal and occipital). Given that blood absorbs NIR light more strongly than the surrounding biological tissue, a hemorrhage is indicated when the comparison between left and right reveal one side is much lower than the other. These devices are typically used for initial screening when injury is suspected in order to recommend further imaging or monitoring, though they do not replace a CT or MRI for SDH diagnosis. These devices do not provide any information related to the depth of the bleed and sometimes indicate injury solely due to extracranial bleeding, which would not require immediate neurosurgical intervention. Additionally, they can struggle in the presence of bilateral injury, due to the comparison-based measurement. Nonetheless, these devices showcase the potential for NIRS to provide a portable, noninvasive tool for diagnosing extracranial bleeds.

A large animal model of life-threatening SDH would be beneficial in order to develop new prehospital diagnostic and therapeutic approaches. Here, we report the development and initial characterization of a sheep model of SDH involving infusion of autologous blood into the subdural space with continuous ICP monitoring. Sheep are an appropriate species for such a model because their skull thickness is similar to that of humans (*34*). For prehospital diagnostic and therapeutic approaches, skull thickness plays a critical role in the fidelity of assessments(*35*). In contrast, most prior large animal SDH models have involved pigs (*36*) (*37, 38*) (*39*) , which have a much thicker skull (**Fig. S1**). Furthermore, most prior large-animal SDH models have involved relatively modest SDH volumes and moderate elevations of ICP, not fully mimicking the life-threatening effects of large SDH in humans (*36*) (*37, 38*) (*39*). Sheep have been previously used to model other forms of traumatic brain injury, with good fidelity and reliability (*40*) (*41*) (*42*) (*43*) (*44*).

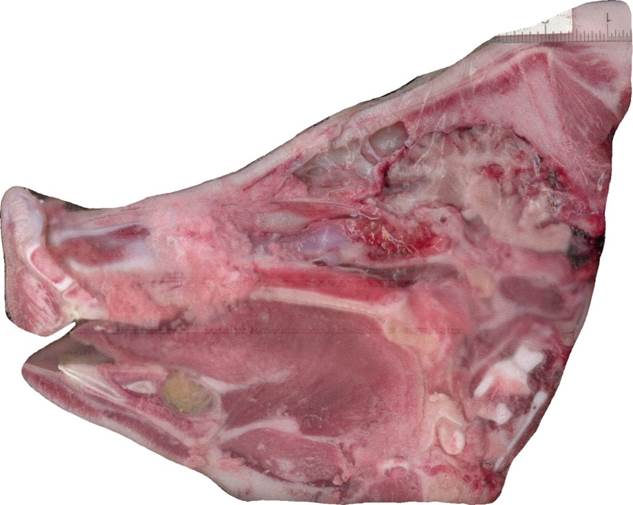

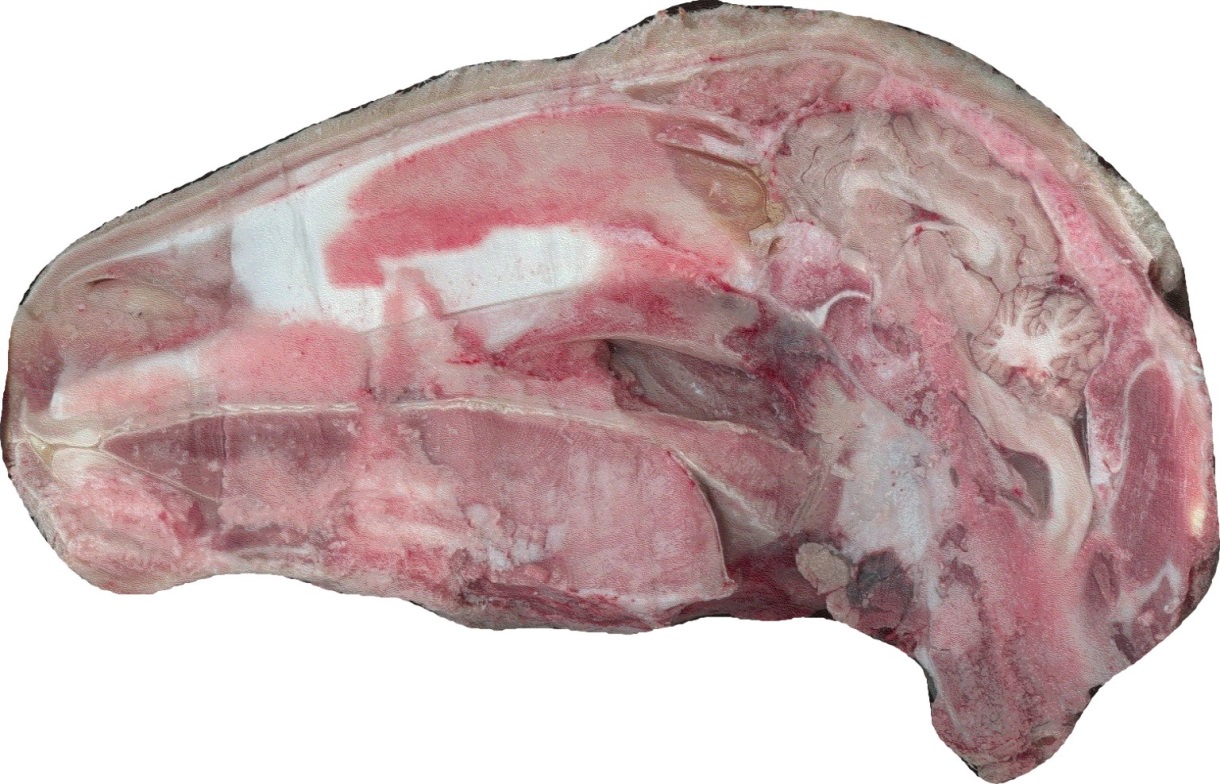

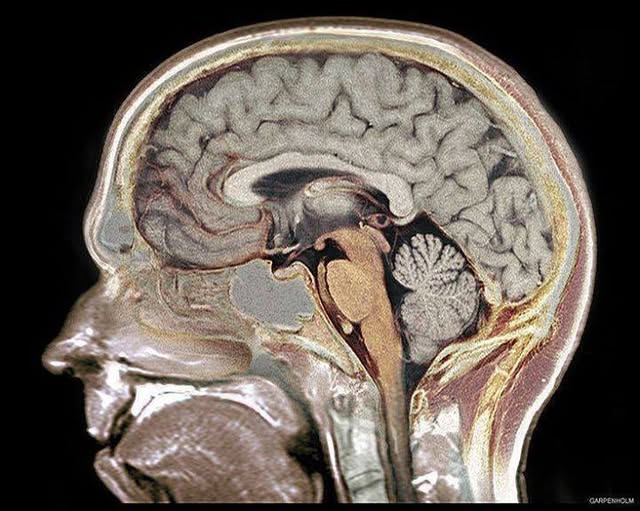

Fig. S1: Sagittal photographs of a human head, a sheep head, and a pig head, showing similar skull thickness in the human and the sheep, but dramatically thicker skull in the pig in the area overlying the brain (green arrows). (Credit: Wikimedia Commons: Garpenholm under Creative Commons and Dr. Adolph Januszkiewicz.

### MATERIALS AND METHODS, SUPPLEMENTARY RESULTS

#### Frequency Domain Multi-Detector Infrared Intracranial Hemorrhage Detection Device (IHDD)

##### IHDD System Overview

The IHDD is a frequency-domain diffuse optical system designed to detect SDH by measuring the optical absorption value in the NIR, since blood is strongly absorbing in this wavelength range. Diffuse optical systems can be implemented using different detection schemes, with trade-offs in levels of complexity, quantitative estimation of optical properties, and equipment size and cost. The simplest detection scheme is known as continuous wave, where the light is continuously delivered to the tissue and the detected light is analyzed to measure intensity fluctuations over time. This is typically used for functional brain imaging (e.g., functional near-infrared spectroscopy or fNIRS) to localize areas of oxygen consumption in the brain during specific tasks. However, the absolute levels of absorption and scattering cannot be measured using this approach. Rather, only temporal variations or spatial comparisons can be measured, which makes it less suitable for diagnostics. Both frequency-domain (FD) and time-domain detection schemes enable absolute measurement of tissue optical properties. FD implementation requires amplitude modulation of the light delivered to the tissue, and the detector is designed to measure the resulting amplitude and phase shift of the light at the detector (**Fig. 1B**). In contrast, time-domain requires a pulsed light source and a detector to resolve the distribution of time of flight for light that travels to the detector. From both the frequency-domain and time-domain data, these measurements are paired with analytic or computational models that describe the diffuse propagation of light through tissue and allow for the estimates of the optical absorption and scattering properties to be calculated. FD is typically less costly and less complex to implement compared to time-domain and has comparable performance in macroscopic optical property estimation. Additionally, the technology is more amenable to a small, portable form-factor. Therefore, a FD design was chosen for the IHDD.

Custom hardware was implemented to modulate the intensity of the light incident on the tissue and to detect the amplitude and phase shift of the light that is collected at the detector (**Fig. S2**). The illumination is achieved with a temperature-controlled laser diode which delivers light to the tissue through a single transmit fiber. The amplitude modulation of the laser is a configurable sine-wave between 100 and 500 MHz. Three detectors (silicon photomultipliers) simultaneously acquire data from three different optical fibers, each spaced an additional centimeter from the transmit fiber (**Fig. 1**). The detected signal is downmixed to an intermediate frequency, digitized and then demodulated in digital signal processing, resulting in the final measurement of amplitude and phase for each source-detector pair (**Fig. S2**). Details of the system architecture and implementation can be found in (*45*), with the notable difference of using a single side-band modulator on the transmit side in order to accurately track global phase of the system, which was needed to properly measure absolute phase delay imparted by the sample. The system built for this effort was designed to allow for portability and maneuverability in the operating room for in vivo testing, while utilizing standard benchtop equipment in order to maintain a degree of flexibility to the hardware to allow for system refinement. Moving forward, the system can be redesigned with the identified performance criteria to build a much smaller system.

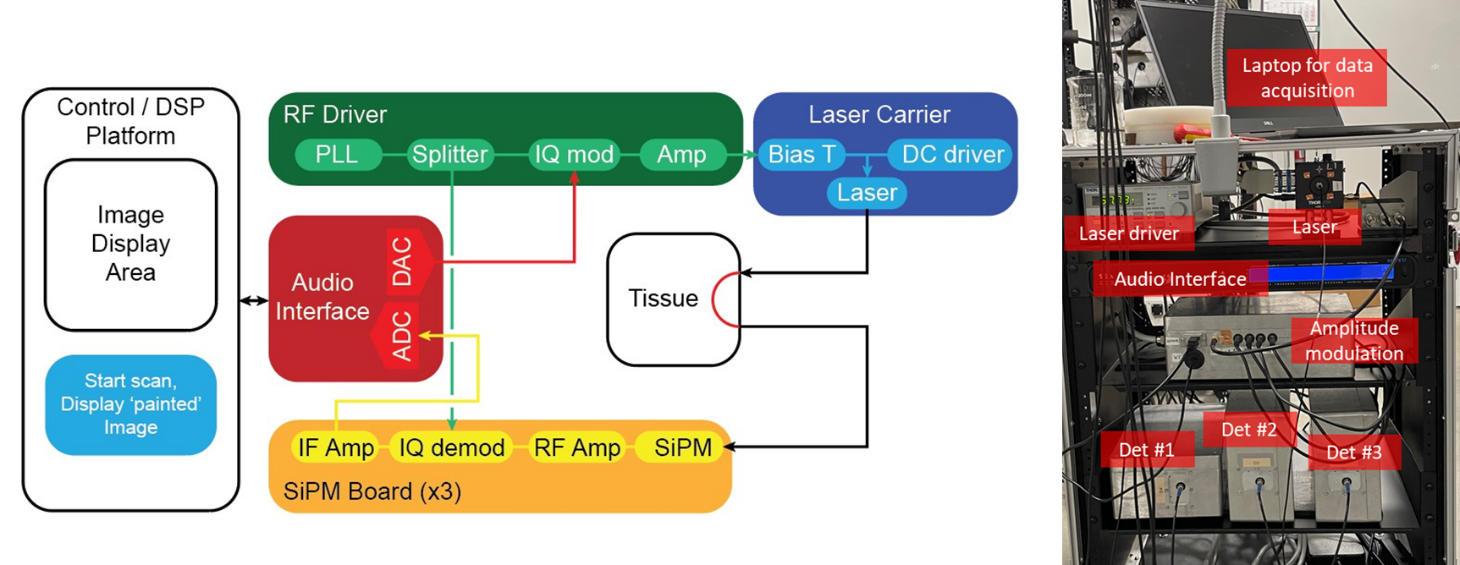

**Fig. S2.** Left: Diagram of the demodulation in digital signal processing. Right: Photo of the benchtop experimental system used for data collection. For convenience, all of the equipment was rack mounted, but each of the components can be readily miniaturized.

The handheld probe used for measurement is tethered to the system via optical fibers and is scanned across the head during data acquisition to measure the entire sample. The probe contains one source fiber and three detection fibers positioned at three different distances (10, 20, 30 mm) from the source fiber. The range of separation distances between the source and detection fiber pair result in some depth selectivity (**Fig. 1A**). The system was designed to acquire from three different detector fibers simultaneously in order to distinguish between extra-cranial and intra-cranial bleeds.

##### Spatial Mapping of IHDD Information:

Position tracking of the handheld probe was incorporated into the data acquisition to provide spatial mapping of the optical data and enable localization of the injury. A position and orientation sensor (Polhemus Patriot system, Colchester, Vermont) was attached to the handheld probe to map the location of each of the receive fibers as the sensor was scanned across the entire head. The data acquisition software (MATLAB, The MathWorks, Inc., Natick MA) populated a spatial map in real-time, showing the location of the mid-point between the source and receive fiber for each of the source-detector pairs (**Fig. 1D**). This provided feedback to the user to ensure that the entire area was sufficiently sampled. Once a scan was complete, all of the data points were used to create a surface map by performing interpolation on the discrete data points. The optical measurements at each of these locations were used to create optical absorption maps (see Data Analysis section).

##### In Vitro IHDD Data Calibration:

To calibrate the system response, a two-step approach was used. First, the optical properties of a homogeneous optical phantom were derived from amplitude and phase measurements at several source-detector separation distances, using a single source-detector pair. Each source-detector separation measurement of amplitude and phase can be referenced to the shortest source-detector separation measurement through complex division, effectively removing system phase lag and system amplitude attenuation. Using the frequency domain standard diffusion equation for a semi-infinite medium with the extrapolated boundary condition (*46*), absorption and reduced scattering parameters can be calculated. These referenced amplitude and phase measurements were fit to the multi-distance standard diffusion equation model through a nonlinear Levenberg–Marquardt least-squares algorithm. This approach was used to characterize an optical phantom fabricated from polydimethylsiloxane doped with black ink and TiO_2_ to control the absorption and scattering characteristics (*47*). Once this homogeneous phantom’s optical properties were characterized, it was then used as the reference phantom to enable the much easier calibration approach during experiments. For this approach, before each sheep scan was collected, the handheld optical probe of the IHDD was used to collect optical data from the reference phantom and then compare it to its theoretical response as estimated using the standard diffusion equation. This difference is the system’s response. The measured phase and amplitudes were adjusted according to this calibration in order to correct for the system response.

#### Sheep Model of Life-Threatening Subdural Hematoma

##### Model Development

The initial model development took place at Thomas D. (TD) Morris Inc. Subsequent testing of the IHDD and IHTD was conducted at Uniformed Services University of the Health Sciences (USUHS). Research was conducted in compliance with DOD and federal statutes/regulations relating to animals and experiments involving animals and adhered to principals stated in *The Guide for the Care and Use of Laboratory Animals,* National Research Council (2011). The study reported herein was performed in animal facilities fully accredited by the American Association for the Accreditation of Laboratory Animal Care. Research was conducted in accordance with the Institutional Care and Use Committees and reviewed by the funding agencies.

Adult female sheep weighing approximately 45-50 kg were purchased from TD Morris at 10-12 months of age. Q-fever testing was performed and verified to be negative prior to purchase. Sheep were housed in groups of 2-4 for at least 7 days prior to experiments. Sheep were provided with a pelleted chow (5508 Rumilab Diet, or equivalent) and alfalfa hay for supplemental roughage and enrichment, ad lib access to water, and environmental enrichment. Light dark cycle was maintained at 12-hour on 12-hour off intervals. Veterinary staff performed daily evaluations of the sheep and verified that they were in good health prior to experiments.

Sheep were prepared for experiments as follows: Sheep were fasted starting 24 hours before the procedure to reduce bloat. Sheep were sedated with 10 mg/kg ketamine plus 0.22 mg/kg xylazine via intramuscular injection in the hind limb under gentle restraint in the home cage. When unconscious, sheep were moved to the operating room and placed on a warming pad in dorsal recumbency. The neck was shaved and a single lumen central jugular venous catheter (Arrowg+ard Blue PSI, Teleflex, Morrisville, NC) was placed percutaneously. Intravenous propofol was then started at a rate of 1.5 mL/min. An additional 0.5 mg/kg of ketamine or 2 mL of propofol were given if needed. A section of abdomen was shaved and a Bovie grounding pad was taped into place. A rumen tube was placed into the stomach to prevent bloat. Sheep were intubated with a 7.5 mm or 8.0 mm cuffed endotracheal tube under direct visualization. Sheep were then turned and placed in sternal recumbency. Blood pressure was monitored using an arterial line placed in the palmar artery. Heart rate and oxygen saturation were assessed continuously using a Mindray Passport 12 (DRE Med, Louisville KY) or portable monitor with an ear clip. End tidal CO_2_ and respiratory rate were assessed continuously via the endotracheal tube. Temperature was assessed every 15 min via a rectal thermometer. Sheep were warmed with additional forced warm air if temperature dropped below 99⁰F. Propofol was infused at 1.25-2.25 mL/min (0.4-0.8 mg/kg) for maintenance of anesthesia. Buprenorphine at 0.06 mg/kg intramuscularly was provided for analgesia and Ancef, 1 g intravenously was administered to reduce risk of infections 30 minutes before intracranial procedures were started.

Two intracranial procedures were performed: placement of an intracranial pressure catheter and placement of a subdural catheter (**Fig. S3**). The sheep was placed in sternal recumbency, the head was shaved to less than 1 mm and washed with saline. A marking pen was used to indicate the anterior and posterior midline of the skull, and the lateral locations for the incisions for burr holes (**Fig. S3A**). Then the skin was prepped with betadine and let dry for 5 minutes. A Bovie electrocautery pen was used to make a 2 cm anterior to posterior skin incision located 4 cm right of midline and 10 cm posterior to the posterior canthus of the eye (**Fig. S3B**). The Bovie was used to clean the periosteum. A 3 mm diamond burr bit attached to a high-speed motorized drill (Micro Medical Electric Spine Drill, model W-MZ-1, OrthoProMed, Middletown DE) was used to create a burr hole down to the dura. The dura was punctured with sharp forceps. A Raumedic pressure transducer (NEUROVENT-P and a MPR-2 datalogger, Raumedic Mills River, NC) was placed through the burr hole into the right frontal cortex at a depth of 1.5 cm and sutured into place with a purse string suture. Baseline intracranial pressure of 4-12 mm Hg with respiratory variation was verified to confirm correct placement. Then, another burr hole was placed contralaterally and widened to 6 mm in an anterior-posterior direction using the high-speed drill (**Fig. S3C**). The dura was punctured, and a 5 French small animal urinary catheter (Mila International Florence, KY) was placed at a gentle angle 2 cm anteriorly into the subdural space. The subdural catheter was glued into place using cyanoacrylate medical grade tissue adhesive (Vetbond 3M, St. Paul MN) (**Fig. S3D**). The subdural hematoma was created by injecting autologous blood drawn from the central venous catheter in 1 mL boluses delivered once per minute (**Fig. S3E-F**). Sheep were monitored for apnea during blood infusions and ventilated manually if apneic for more than 30 seconds. Then, the subdural catheter and intracranial pressure monitor were removed, the glue was removed manually, and the incisions were closed with 2-0 silk sutures (Ethicon, Bridgewater NJ) (**Fig. S3G**).

When required, subgaleal hematoma was created by injecting 2 mL of autologous blood into the subgaleal space using at 19-gauge needle in a fan-shaped distribution (**Fig. S6H**).

For neurological evaluations, the sheep were awakened from anesthesia. For pain, they were given buprenorphine at 0.06 mg/kg intramuscularly every 6 hours.

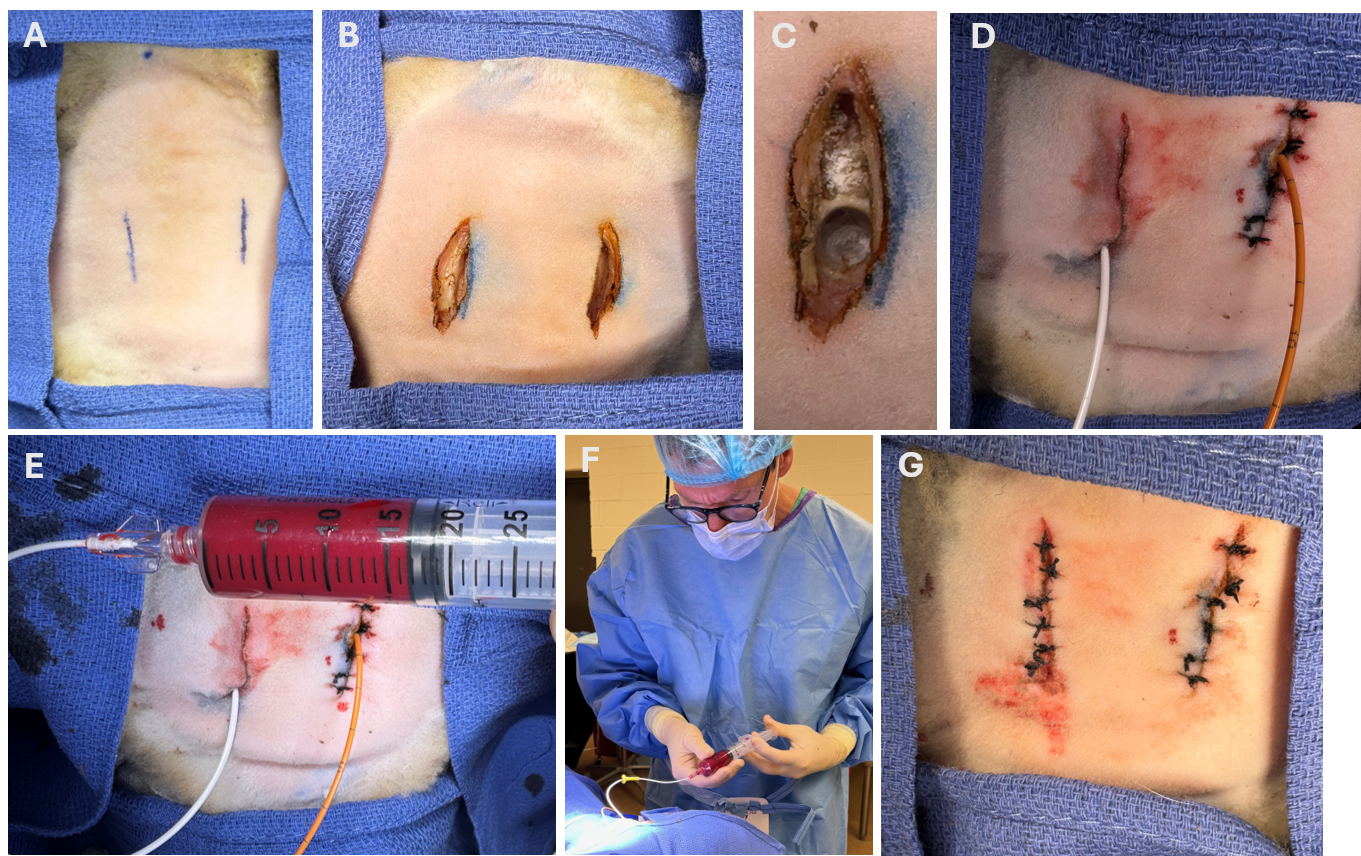

Fig. S3. Creation of the sheep model of life-threatening subdural hematoma including placement of an intracranial pressure catheter and placement of a subdural catheter. **A.** Sheep head shaved, prepped, draped, and marked for location of incisions. **B.** Skin and soft tissue incisions made with Bovie electrocautery. **C.** Burr hole through the skull to dura. **D.** Insertion of intracranial pressure monitor (orange catheter- right side) and subdural catheter (white catheter-left side). **E-F.** Infusion of autologous blood from a 30-cc syringe through the subdural catheter. **G. C**atheters removed and incisions closed with interrupted sutures.

##### Neurological Evaluations:

During the initial model development neurological evaluations were performed twice daily for 72 hours by a board-certified veterinarian and veterinary technician. The evaluations included a 24 point scoring (**Fig. S7**) involving mental status (coma, stupor, lethargy, awake); seizures (present, absent); pupil reflex (unreactive, sluggish, normal); corneal reflex (absent, present); oculovestibular reflex (absent, present); swallowing reflex (absent, present); deep tendon reflexes (absent, hyperactive, normal); step reflex (absent, present); righting (absent, present); standing (unable, abnormal posture, stands with difficulty, normal); coordination (no attempt to walk, attempts to walk but cannot, walks abnormally, walks normally); withdrawal to pain (none, unilateral, bilateral); vision (no reaction, startles to face touch, reacts to approaching object normally). Taken together, these evaluations were used to form an overall assessment of impairment as mild, moderate, or severe.

During the randomized controlled trial of the IHTD (see below) expanded 32-point neurological evaluations were performed because of veterinarian preference (**Fig. S15**).

Neurological status assessments indicated substantial deficits on day 1 and day 2, with gradual recovery in most surviving sheep on days 3 and 4 (**Table S1**).

| **Neurological Score** | | | | | | | |
| --- | --- | --- | --- | --- | --- | --- | --- |
| **Subdural blood infusion (mL)** | **Day** | | | | | | |
|  | **1** | | **2** | | **3** | | **4** |
|  | **a.m.** | **p.m.** | **a.m.** | **p.m.** | **a.m.** | **p.m.** | **a.m.** |
| **10** | 24  N = 3 | 21.3±1.5  N = 3 | 22.3 ± 1.5  N = 3 | 23.0±1.0  N = 3 | 23.7±0.6  N = 3 | 24.0±0.0  N = 3 | 24.0±0.0  N = 3 |
| **12** | 24  N = 6 | 19.0 ± 5.7  N = 6 | 21.7 ± 4.8  N = 6 | 23.8 ± 0.4  N = 5 | 23.8 ± 0.4  N = 5 | 24.0±0.0  N = 5 | 24.0±0.0  N = 5 |
| **15** | 24  N = 6 | 17.0 ± 4.4  N= 5 | 20.0 ± 4.2  N= 5 | 22.7 ± 1.5  N= 3 | 23.7 ± 0.6  N= 3 | 23.7 ± 0.6  N= 3 | 24.0±0.0  N = 3 |
| **20** | 24  N = 3 | N/A | N/A | N/A | N/A | N/A | N/A |

Table S1**:** Neurological assessment for surviving sheep over time after SDH. Neurological scores were based on evaluations by trained Veterinarians using the 24-point scale, where higher scores represent better function. Values represent the mean ± standard deviation of the scores. Number of surviving animals per group is also noted. The a.m. score on the first day is pre-surgery.

##### Euthanasia and Post-Mortem Examination:

For euthanasia, sheep were administered 150 mg/kg intravenous Euthasol (390 mg pentobarbital sodium and 50 mg phenytoin sodium per mL). After confirmation of death, necropsy was performed immediately. The subgaleal space was exposed and the location of any subgaleal hematomas were photographed (**Fig. S6**). Then the skull was opened and the location of any subdural hematomas were photographed. Finally, the brain was extracted and sliced into 1 cm coronal slabs. Portions of the coronal slabs corresponding to left and right parietal cortex, left and right hippocampus plus perihippocampal cortex, cerebellum, and medulla were cut and placed in neutral buffered formalin fixative. The rest of the coronal slabs were placed on dry ice, typically within 50 minutes of euthanasia, and then transferred to a -80 ⁰C freezer.

Post-mortem CT scans of the head were performed instead of rapid necropsy in a subset of the sheep. The CT scans were performed using a Philips Brilliance CT Big Bore scanner with energy 120 kV, 215 mA, helical rotation with 0.44 pitch, 10.5 mm/s table speed, 0.5 s rotation time, 12 mm collimation reconstructed into 32 slices with approximately 1 mm isotropic spatial resolution.

##### ICP Data Extraction and Analysis:

Data from the Raumedic intracranial pressure monitor were exported as .csv files and then analyzed using Microsoft Excel 2019. The time data were cleaned by deleting the date information using the ‘Find/Replace’ function (e.g. Find ‘**2025 ’, Replace ‘ ‘), then subtracting the initial start time in ‘Number’ format) to yield serial time stamps. The intracranial pressure data were cleaned by deleting the initial headers before the semicolon using the ‘Find/Replace’ function (Find: ‘**;’, Replace ‘ ‘). The data were plotted in GraphPad Prism 10.5.0.

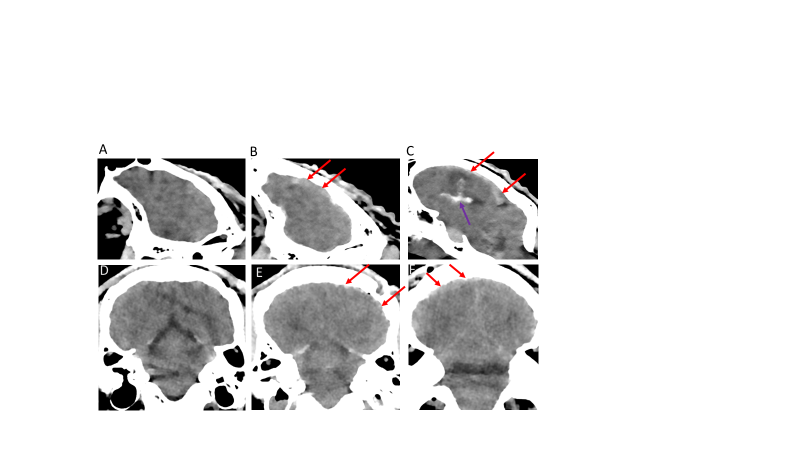

Fig. S4: **A-C** Post-mortem parasagittal CT scans. **A.** sham **B.** SDH **C**. SDH + intraparenchymal and intraventricular hemorrhage. **D-F** coronal CT scans. **D.** Sham. **E.** patchy SDH **F.** SDH crossing midline.

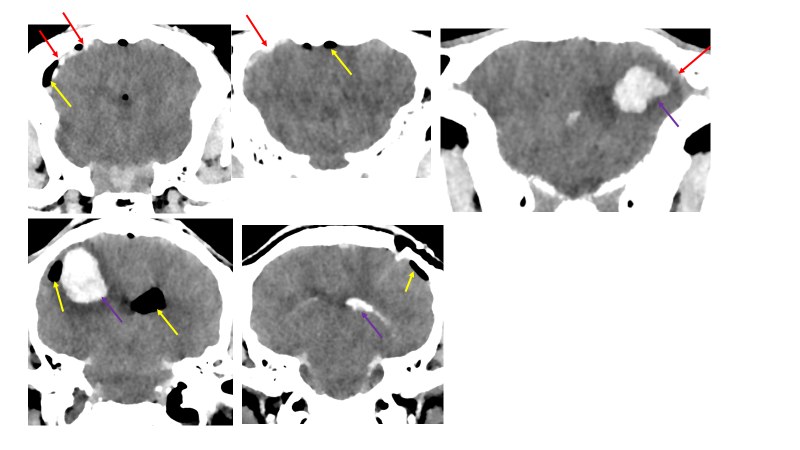

Fig. S5 Additional post-mortem CT scans demonstrating subdural hematoma (red arrows), pneumocephalus (yellow), and intraparenchymal or intraventricular hemorrhages (purple).

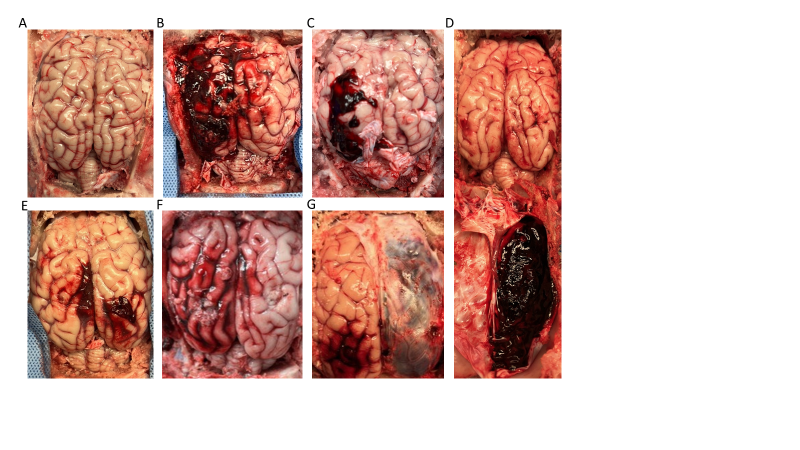

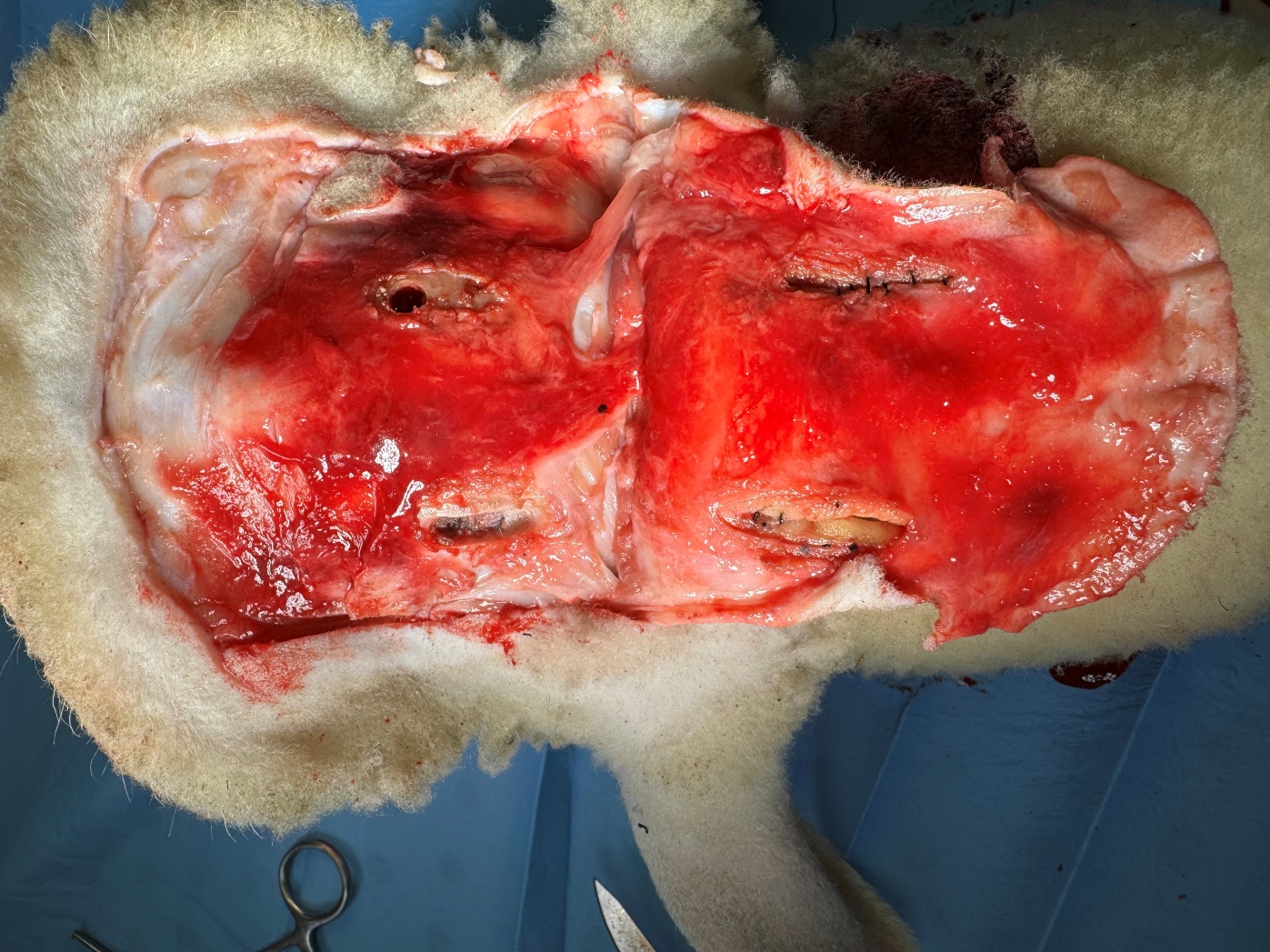

**H**

Fig. S6**.** Additional post-mortem photographs. **A.** Sham procedure with no blood injected. **B.** Left subdural hematoma with some contralateral blood. **C.** Smaller left subdural hematoma; additional blood was found in the basal cisterns and intraventricularly. **D.** Right subdural hematoma with blood adherent to reflected dura. **E.** Sham procedure with small amounts of blood collection around the site of the intracranial pressure catheter and subdural catheter. **F.** Left subdural and subarachnoid hemorrhage with some contralateral blood**. G.** Right subdural hematoma shown through the intact dura. **H.** Bilateral subgaleal blood on the surface of the skull after reflection of the scalp.

| Mental status | Coma, no responsiveness | |
| --- | --- | --- |
|  | Stupor, responsive to vigorous stimulation only with posturing | |
|  | Lethargy, drowsiness or delirium | |
|  | Awake | |
| Cranial nerves | Pupil reflex | |
|  |  | Unreactive (dilated and not constricting to light) |
|  |  | Sluggish (dilated and slow to constrict to light) |
|  |  | Normal |
|  | Corneal reflex | |
|  |  | Absent |
|  |  | Present |
|  | Oculovestibular reflex | |
|  |  | Absent |
|  |  | Present |
| Sucking Reflex | Absent | |
|  | Present | |
| Swallowing Reflex | Absent | |
|  | Present | |
| Reflexes | Deep tendon reflex | |
|  |  | Absent |
|  |  | Hyper |
|  |  | Normal |
|  | Step reflex | |
|  |  | Absent |
|  |  | Normal |
|  | Righting | |
|  |  | Absent |
|  |  | Normal |
| Motor: Global | Standing | |
|  |  | Unable to stand |
|  |  | Bears weight with abnormal posture |
|  |  | Gets to standing with difficulty |
|  |  | Stands normally |
|  | Coordination |  |
|  |  | No attempt to walk |
|  |  | Attempts to walk but cannot |
|  |  | Walks but fails |
|  |  | Walks normally |
| Sensorimotor | Noxious stimuli response |  |
|  |  | Does not respond to pain |
|  |  | Withdraws from pain unilateral (note side) |
|  |  | Withdraws from pain bilateral |
|  | Vision |  |
|  |  | No reaction, startles to face touch bilateral |
|  |  | Startles to face touch unilateral (note side) |
|  |  | Reacts to approaching object; does not startle to face touch |

Fig. S7. Exemplar sheep neurological assessment using the 24-point scale.

#### **Additional Model Characterization**

Infusion of autologous blood into the subdural space of adult sheep resulted in nonlinear increases in intracranial pressure (**Fig. S8**). Initial blood infusions resulted in relatively modest increases in pressure, whereas subsequent blood infusions resulted in larger increases in pressure (**Fig. S8A**). This was interpreted as consistent with loss of intracranial compliance and compensatory reserve. High time resolution recording (every 10 msec) revealed several features of the ICP dynamics: First, towards the end of the blood infusions, when ICP was very high, there were additional sharp spikes in ICP related to each mL of blood infused. These ICP spikes relaxed in two phases, one which decayed over 100’s of milliseconds (**Fig. S8B**), and the slower phase which relaxed over minutes. Second, there was clear variability with heart rate and respiration, and the amplitude of the ICP variability increased at higher pressures (**Fig. S8C-D**). Some experiments showed more irregular changes in ICP, with initial rapid rise in ICP, a relative plateau phase, and then further increases in ICP. There was substantial variability in the extent to which ICP rose after infusion of a constant volume (15 mL) of blood into the subdural space (**Fig. 2F**). Baseline ICP was 12.2 ± 4.6 mm Hg (mean, SD, n=18). After 15 mL of blood infusion, ICP increased significantly to 114 ± 42 mm Hg (repeat measures ANOVA overall F_1.465, 24.90_ = 92.97, p<0.0001, Tukey test p<0.0001 vs baseline). At 5 minutes following blood infusion, ICP had decreased to 85 ± 30 mm Hg, a 24% decrease (p=0.004 vs. after 15 mL of blood infusion). During model development, ICP measurements were terminated after 5 minutes.

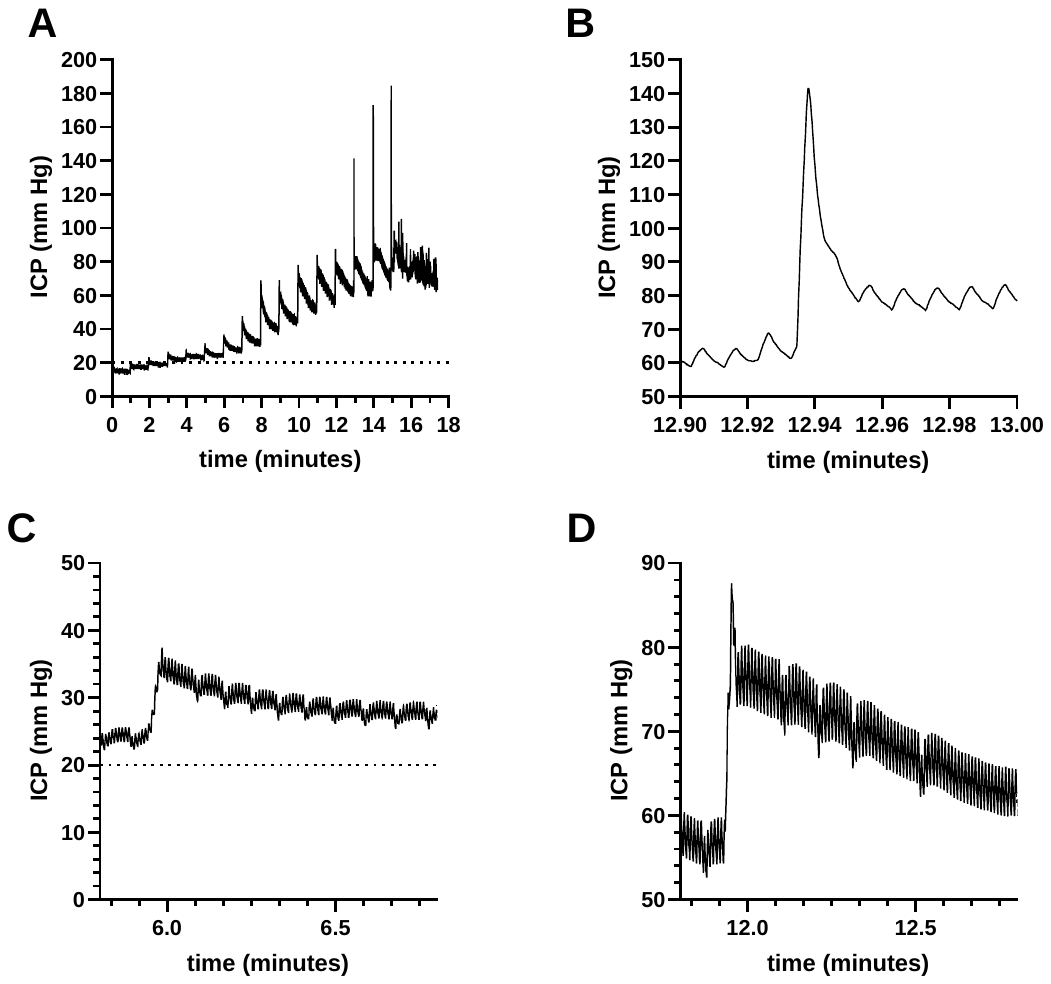

Figure S8: Elevated intracranial pressure (ICP) related to infusion of autologous blood into the subdural space. **A.** Exemplar time course of ICP showing elevations in pressure at the time that each mL was infused. **B.** High time resolution measurements (10 msec sampling) showing rapid ICP spike superimposed on cardiac-related fluctuations. **C.** Higher time resolution ICP just after minute 6 with moderate ICP elevation with both respiratory and cardiac-related fluctuations apparent. **D.** Higher time resolution ICP measurements just after minute 12 with severely elevated ICP, larger amplitude respiratory and cardiac-related fluctuations, and irregular respirations.

There were irregular elevations in blood pressure and variable heart rate responses during subdural infusion of 15 mL blood, consistent with Cushing response (**Fig. S9-S10**). In 20 sheep with 15 mL of subdural blood infusion, arterial catheters were placed for measurement of blood pressure. Measurements were recorded once per minute just prior to infusion of each mL of blood into the subdural space. Mean arterial blood pressure increased from 111 (SD 8.4) mm Hg at baseline to 129 (SD 39.6) after 15 mL of subdural blood infusion. Of note, although most sheep had hypertensive responses, this was not universal. Pulse pressure increased from 26 (SD 9.1) mm Hg at baseline to 43 (SD 33.9) after 15 mL of subdural blood infusion. Heart rate decreased from 133 (SD 24.7) beats per minute at baseline to 112 (SD 52.3) after 15 mL of subdural blood infusion. Of note, although most sheep had bradycardic responses, 7/18 had pronounced tachycardic response. Thus, the intracranial pressure increases induced by subdural blood infusion generally had the expected effect on autonomic function in the sheep, even under propofol anesthesia, though there was considerable variability from animal to animal.

.
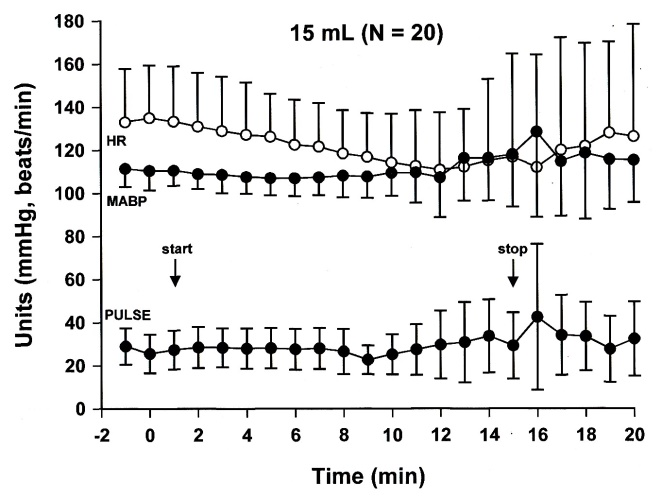

Figure S9. Hemodynamics during SDH induction, including heart rate (HR), Mean arterial blood pressure (MABP), and pulse pressure (systolic – diastolic) from the start if SDH induction at minute 1, to the end at minute 15. (means and standard deviations)

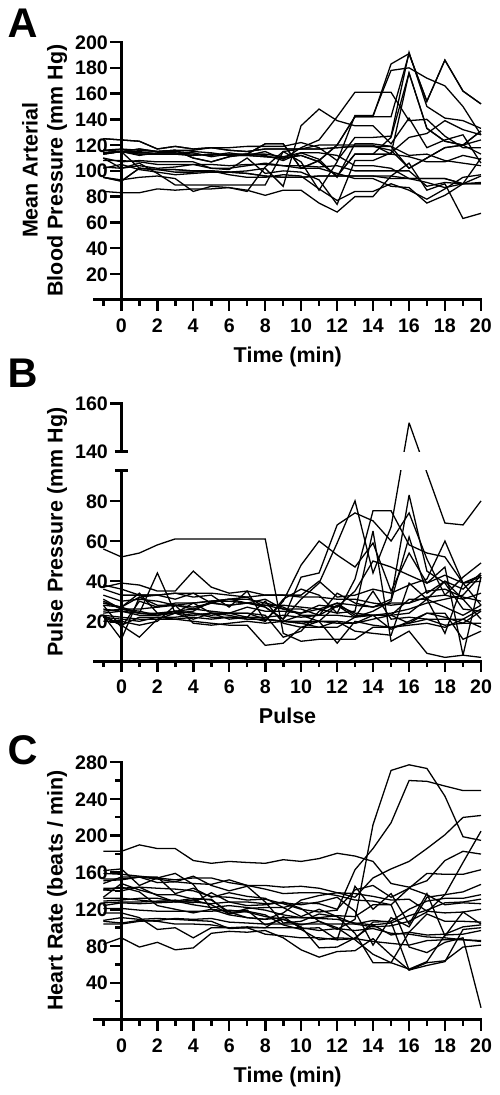

Figure S10: Individual variability in cardiovascular response to infusion of 15 ml of autologous blood into the subdural space in adult sheep measured via arterial catheters. **A.** Mean arterial blood pressure. **B.** Pulse pressure (systolic – diastolic) **C.** Heart rate.

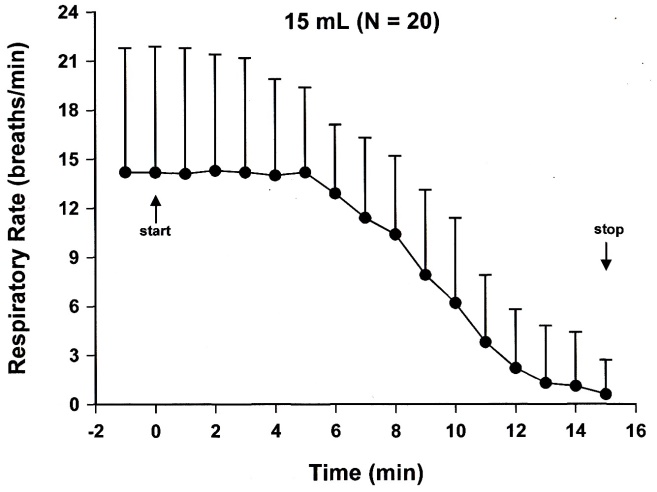
Respiratory rate declined with infusion of blood into the subdural space (**Fig S11**). When sheep demonstrated apnea or hypopnea as judged by the veterinary team, they were placed on assist control ventilation at a rate of 10 breaths per minute and positive end expiratory pressure of 5 mm Hg. Sheep were given room air to breathe, supplemented with oxygen as needed to keep pulse oximetry saturation above 90% as measured on the ear. On average, reduction in respiratory rate occurred in a graded fashion, but in each individual sheep, the onset of hypopnea was more abrupt, with substantial variability from sheep to sheep in when the reduction began.

Figure S11. Respiration rate during SDH induction (means and standard deviations)

Reduction in respiratory rate typically occurred after ICP reached a threshold of 77 mm Hg (**Table S2**). At this ICP, MAP was 110 mm Hg on average, and cerebral perfusion pressure was approximately 33. Apnea typically occurred when ICP reached an average of 93 mm Hg. At this ICP, MAP was still approximately 110 mm Hg on average and cerebral perfusion pressure was approximately 17.

| **Respiration (N = 20)** | **ICP ± s.d. (mmHg)** | **Injected Blood Volume** |
| --- | --- | --- |
| **Normal (threshold)** | 77.6 ± 16.5 | ~ 10^th^ mL |
| **>50% RR Decrease** | 84.6 ± 19.6 | ~ 11^th^ mL |
| **Apnea** | 93.0 ± 28.3 | ~ 12^th^ mL |

Table S2. Transition from normal breathing to apnea occurred from approximately the 10^th^ to 12^th^ mL of subdural infused blood.

There was variability in the localization of blood apparent in the intracranial space on post-mortem CT and at necropsy. In most cases, ipsilateral subdural blood was present as expected (**Fig. S4-S6**), but there were also incidences of contralateral subdural hematoma, intraparenchymal hemorrhage, intraventricular hemorrhage, and blood in the basal cisterns**.** Subdural hematoma, intraparenchymal hemorrhage and intraventricular hemorrhage were readily detected. Subdural hematoma was less salient on CT in the sheep model than is often observed in human patients, in part likely due to the small size of the sheep intracranial vault.

Similarly, there was vaiability in the extent and localization of subdural blood apparent on direct post-mortem examination (**Fig. S6**). There were examples with some contralateral bood, and occassional bleeding around the stie of the intracranial pressure catheter and subdural catheter insertion.

Taken together, these findings indicated that infusion of 15 mL of autologous blood into the subdural space of adult sheep resulted in features comparable to severe life-threatening SDH in humans. However, there was variability in the model. Furthermore, it is possible that the Cushing response may have been blunted by propofol anesthesia.

#### In Vivo IHDD Data Collection:

##### Expert Users and Medical Providers

IHDD data were collected by two different user groups prior to injury as well as after the first randomized injury and the subsequent randomized subgaleal blood infusion. The first group was made up of experts who built the system and had an intricate knowledge of the physics of the sensor and hardware used to build the system. The second user group was made up of users with medical training similar to those who would ultimately be using this system for medical diagnoses. Written IHDD training was provided to the medical providers and hands-on practice scans were performed with guidance from the IHDD engineers before they began collecting experimental data. For each experiment, one expert user and two trained medical providers collected IHDD data and Infrascanner data during the preinjury state, after the first randomized injury and after the subsequent randomized subgaleal blood infusion. The trained medical providers were blinded to the injury status. These two different user groups allow for important comparisons on data quality and offer insight into the usability and interpretability by the target end-users.

##### Infrascanner 2000

In addition to the scans performed with the IHDD, data was also collected with the Infrascanner 2000, a commercially available device marketed for detecting intracranial bleeding, in order to compare and contrast performance between the two devices. The Infrascanner is designed to collect data at distinct areas on the human head and compare the same locations on the left and right side of the head to determine if bleeding is present. For these experiments in sheep, the scan pattern was adjusted to adapt to the much smaller surface area of the cranial compartment. Two pairs of measurements were made, each with the source-detector line oriented parallel to midline; the first pair of measurements were closely spaced (~1cm) on either side of midline and the second pair probed a larger distance from midline (~2.5cm). Positive detection in a hemisphere was made if the difference in optical density for either measurement pair was higher than the threshold provided in the device manual. For both the IHDD and the Infrascanner, data acquisition times typically took 3-4 minutes, with a range of 2-5 minutes.

##### Multi-frequency IHDD data

During all experiments, the expert users collected data with multiple modulation frequencies in order to assess if these additional data improved the ability to discriminate between the different injury cases. Data was collected at 100, 200, 300, 400 and 500 MHz. These data were utilized for different approaches to injury classification as well as correlation analysis between the different modulation frequencies. While a modest improvement was observed in most cases, this improvement was attributed to having more repeat measurements at the same locations, rather than unique information measured from the different modulation frequencies. Given that the injury diagnostic accuracy was not impacted very significantly from this additional system complexity, only the 200 MHz dataset was carried forward for injury classification quantification.

##### IHDD Data Analysis

While the raw amplitude and phase contain a wealth of information, it also contains several confounding factors. Any change in optical coupling, cable length, digitizer phase offset, or detector efficiency will also affect the amplitude and phase of the signal collected. In order to separate out the system component effects from the sample, a system calibration was performed using an optical phantom with known optical properties. Moreover, the calibrated amplitude and phase are nonlinearly coupled to physiological states of interest (e.g., change in blood volume). We implemented optical data analysis to separate the contributions of optical absorption from scattering so as to more directly correlate the changes in physiology to changes in signal collection. Using the standard diffusion equation theoretical model for a semi-infinite medium with the extrapolated boundary condition (*46*), the optical properties were calculated at each source-detector pair over time using the known source-detector separations of each nearest neighbor, the modulation frequency of the source, and the calibrated amplitude and phase measurements (*48*). These parameters were inputs to an iterative solver using the nonlinear Levenberg–Marquardt least-squares algorithm.

For improved visual interpretation and inter-scan comparison, the position and optical data were resampled to a regular grid (**Fig. 1D**). First, the 3D position data was projected onto the XY plane, then a linear interpolant of 1 mm spacing resampling each metric of interest, e.g., absorption, amplitude, and phase was created. Hence, for a given sheep, there is a rich data set including injury state, source-detector separation, optical metrics, and position.

##### Support Vector Machine Classifier

Automated analyses were performed in order to aid in determining if a strong absorption signal is a life-threatening SDH versus non-life-threatening subgaleal blood. A machine-learning based model was created to estimate the likelihood of SDH, subgaleal hemorrhage, or both at each location using the phase shift and amplitude attenuation of the infrared light from all 3 detectors. Classification was performed using an error-correcting output codes multi-class support vector machine with one-vs-one coding and cubic polynomial kernel base learners. The classifier was trained using labeled data from 14 subjects. The classifier was subsequently evaluated on previously unseen data from 17 subjects. Some of the data came from the model development phase and some from the randomized blinded study. The classifier used amplitude, phase, and absorption from all three channels to discriminate among the 4 classes: (1) no hemorrhage present, (2) SDH only, (3) subgaleal blood only, and (4) both subgaleal blood and SGH present together. Classification was run on individual pixels within each scan. Additional criteria of continuity and count were leveraged to determine the overall injury diagnosis for the left and right hemispheres. Any areas of pixels with contiguous SDH labels that were smaller than 4 total pixels were removed. Then, the total number of pixels with SDH or Both labels were summed together and compared to the number of subgaleal blood pixels. If the former group was larger than the latter group and of a minimum size set by a threshold, that hemisphere was labeled as SDH injury. If the maximum area of the subgaleal blood pixels was 1.5x greater than the SDH and at least the size of the threshold, it was ruled subgaleal blood.

We assessed the possibility that the support vector machine classifier could provide greater accuracy than the medical providers with 30 minutes of training each. The model output included a classifier which assigned each pixel to one of 4 categories: none, subgaleal hemorrhage, SDH, or both. The classifier was trained based on ground-truth results from 14 sheep and then tested on data from 17 additional unseen sheep. The classifier results were compared with the ground truth for these 17 additional sheep.

Overall, the support vector machine classifier provided similar accuracy for trained medical providers to direct diagnoses made based on visualization of the attenuation from all 3 detectors (**Table S3**). Data from expert users was somewhat more accurate, but the expert users were not blinded, so this result should be interpreted with caution.

| **Device** | **User Group** | **Sensitivity (95% CI)** | **Specificity (95% CI)** |
| --- | --- | --- | --- |
| IHDD | Expert Users | 75% (53-90%) | 66% (51-79%) |
| IHDD | Medical Providers | 69% (56-82%) | 51% (38-63%) |
| Infrascanner | Expert Users | 45% (24-68%) | 76% (61-88%) |
| Infrascanner | Medical Providers | 52% (37-67%) | 54% (40-67%) |

**Table S3.** Sensitivity and specificity of IHDD data using support vector machine classification for expert users and trained medical providers, along with Infrascanner 2000 data from the same experimental subjects.

##### Infrascanner Performance in Sheep Model

Measurements with the Infrascanner were collected at each stage of the injury, either directly before or after the IHDD scans in order to assess Infrascanner performance for each injury. The scan pattern was adapted for the sheep head. The raw optical density (OD) values recorded by the scanner were imported for analysis and diagnosis was made using the OD threshold given in the Infrascanner manual. The sensitivity and specificity calculated from the diagnoses were tabulated from the scans comparing opposite hemispheres (**Table S3**). The sensitivity across user groups had a larger overlap than the specificity between user groups, which may indicate that false positives were more prevalent for trained users compared to expert users. Additional training and experience with the device could help to get consistent performance of the device across all users.

There was less variability between users in accuracy for the new infrared hemorrhage detection device compared with the Infrascanner 2000. Range of accuracy for the new infrared hemorrhage detection device was 0.56 to 0.75, with a standard deviation of 0.069. Range of accuracy for the Infrascanner 2000 was 0.43 to 0.79, with a standard deviation of 0.14. The difference in variance was not statistically significant (F-test, p=0.14)

In four additional sheep, the same users performed assessments using both devices, to determine whether there might be synergistic information obtained. Instead, accuracy was even worse than for either device in isolation. For the simple scenario of subdural vs sham, sensitivity was 0.48 and specificity was 0.57, with an overall accuracy of 0.52. For the more complex scenario including subgaleal blood, sensitivity was 0.76 and specificity was 0.29, with an overall accuracy of 0.53. Thus, using both devices is not likely to be beneficial.

#### Intracranial Hemorrhage Treatment Device (IHTD)

##### Concept and Requirements

The overall concept for the IHTD involved a single device with all of the functions required to create a burr hole in the skull of a patient with life-threatening subdural hematoma, incise the dura, and allow access to the subdural space. Our military medical advisors recommended a single device, rather than a kit containing multiple parts which could get lost or be difficult to find. They also advised us that the smaller and lighter the device could be made, the more likely it would be to be routinely used by military medical providers who have tight constraints on equipment and supplies. A third piece of advice was to be mindful of the risk of infection given the possibility of needing to perform prolonged field care for up to 72 hours in an austere environment. In austere environments, culture and sensitivity-based antibiotic treatment, as would occur in a fully staffed hospital, would not be possible; our advisors noted that a high risk of death from meningitis or encephalitis due to an open burr hole was a concern.

Based on this advice, we formulated several requirements:
1) A single closed device with multiple functional units. The functional units should be designed to be moved into place over the area where the burr hole would be created. We selected a disk design that allowed the functional units to rotate into place and lock in, then be freed to rotate out using a release button (**Fig. S12**).

2) A way to fasten the device to the patient for up to 72 hours, including possible transport with the device attached. Attempts to use adhesives (**Fig. S12**) did not meet operational requirements, and a system involving 3 bone screws was adopted for subsequent designs (**Fig. S13A, S14A**). We considered systems involving powered screwdrivers but ultimately decided that hand-tightened screws would be lighter and less likely to break.

3) A tight seal against the skin to reduce the risk of infection in an austere environment. This was accomplished using a closed-cell foam rubber gasket on the bottom face of the device (**Fig. S13A, Fig. S14B**). Once a life-threatening subdural hematoma was identified, the plan would be to shave the relevant region of the head, clean the skin to remove visible contamination with blood, dirt, or other foreign material, decontaminate the skin with a surgical prep such as betadine or alcohol, and then affix the treatment device with a tight seal. We recognize that it may not be possible to keep the treatment site fully sterile, but the goal was to reduce the risk of life-threatening meningitis or encephalitis in a prolonged field care setting.

4) A method to penetrate the scalp and expose the skull bone. In early testing, we recognized that the drills intended to make a burr hole in the skull bone would slip off of the scalp and not penetrate well. We also recognized that profuse scalp bleeding caused by incising the scalp with a blade could rapidly reduce visibility in the area intended for the burr hole. For these reasons, we chose an electrocautery system that both cuts through the scalp and staunches bleeding. To implement this plan, a punch biopsy blade was affixed to a tool made of thermally conductive metal. An electrical resistive heating element and thermocouple were embedded in the tool and connected to an external battery and monitoring circuit (**Fig. S13B, Fig. S14C**). The circuit included a temperature feedback controller which adjusted the current to achieve a temperature of 232 ⁰C, which was found to be appropriate for rapid cutting and coagulation of scalp tissue.

5) A method to remove the plug of scalp tissue from the site where the burr hole would be made. In early testing, we found that the scalp tissue variably interfered with subsequent visualization if left in place after electrocautery. Therefore, we designed a sharpened coil intended to hook the plug of scalp tissue and remove it from the field (**Fig. S13C, Fig. S14D**).

6) A method to create a burr hole without damaging the underlying brain in case the user made an error in localizing the subdural hematoma. We selected a hand cranked drill rather than a powered drill, to reduce weight and increase robustness (**Fig. S13D-E**). The drill system was designed using a threaded stem and a clutch to allow the drill to advance while limiting the axial force that is able to be applied. This force limit reduces the torque required for drilling and prevents the drill from advancing. A hard stop contacting the outer surface of the skull also prevented excessive penetration. The drill bit was approximately 1 cm in diameter.

7) A method to incise the dura. The drill also cut through the dura in some cases, but this was highly variable and in other cases the dura remained intact (**Fig. S14E**) and intracranial pressure did not drop until the dura was incised (**Fig. S16**). We therefore added a deployable blade attached to the drill bit that could be used to cut the dura after the burr hole had been created **(Fig. S14G**). During the burr hole drilling, the blade remained tight against the drill bit and did not protrude so that it would not get broken by the bony skull. Once the burr hole had been created, the blade was deployed forward by 1 mm past the drill tip. One more 360-degree rotation of the drill assembly with the blade deployed was sufficient to cut the dura (**Fig. S14F**).

8) A method to provide access to the intracranial space in order to remove as much of the subdural hematoma as possible. We recognized that acute subdural blood would clot rapidly in the subdural space, and simple aspiration would not be effective. Therefore, we designed the device to be compatible with the NICO Myriad morcellator, which has been previously demonstrated to be useful for treatment of lobar intracranial hemorrhages in the operating room leading to improved neurological outcomes (*49*). We added small access port (**Fig. S13F**) with dimensions designed to allow the handpiece of the NICO Myriad morcellator to pass through it (**Fig. S14H**). The access port was made as small as possible to reduce the risk of infection. In the future, the access port could be made of a material that re-seals after the hand piece has been removed for even better infection control.

The prototypes were constructed at the Johns Hopkins Applied Physics Laboratory from a combination of off-the-shelf parts and custom fabricated components.

A US Patent, US 12,150,655B2, Brody et al. Date: Nov. 26, 2024, entitled “AUTOMATED CRANIAL BURR HOLE DEVICE AND METHOD” has been granted.

Fig. S12: The initial design of the IHTD intended for proof-of-concept testing in a non-living human scalp and skull surrogate. **A.** Photograph of the device on the bench top, with red and black electrical cord and yellow connector for attachment to power source for electrocautery. **B.** Size scale on human head surrogate. **C.** Proof-of-concept components including adhesive and sealing base stage and electrocautery, drill, and treatment access port on a rotating upper stage.

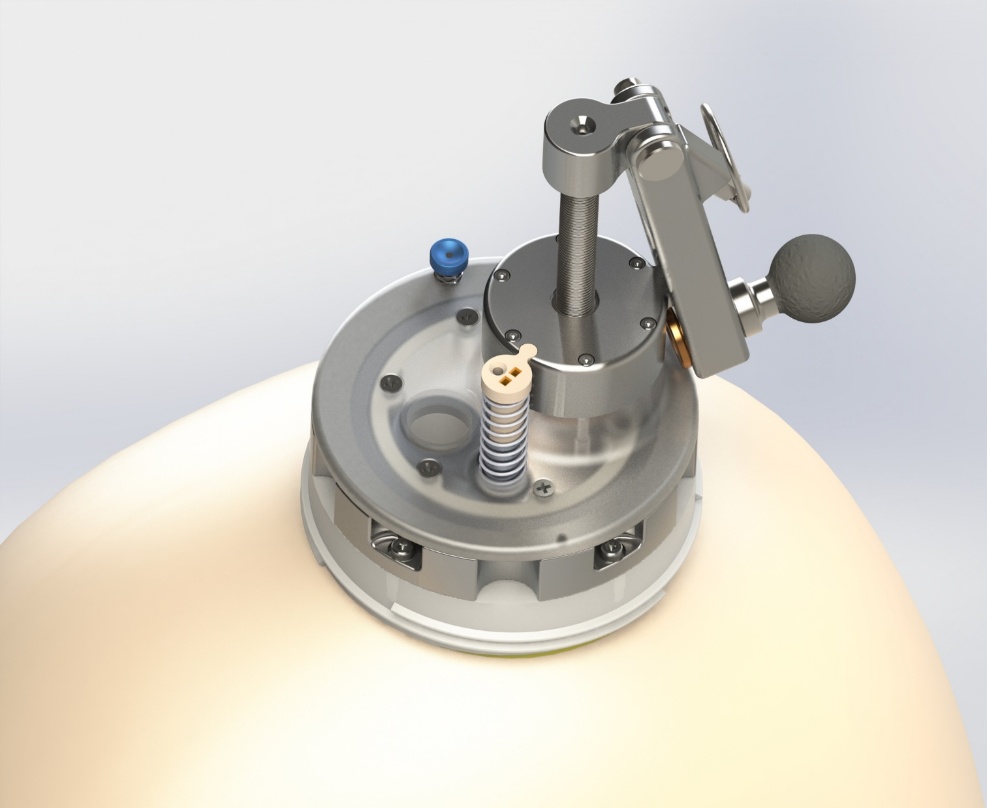

Manual drilling mechanism

Release button to allow lid rotation

Treatment access port

Electrocautery plunger

Adhesive and sealing stage

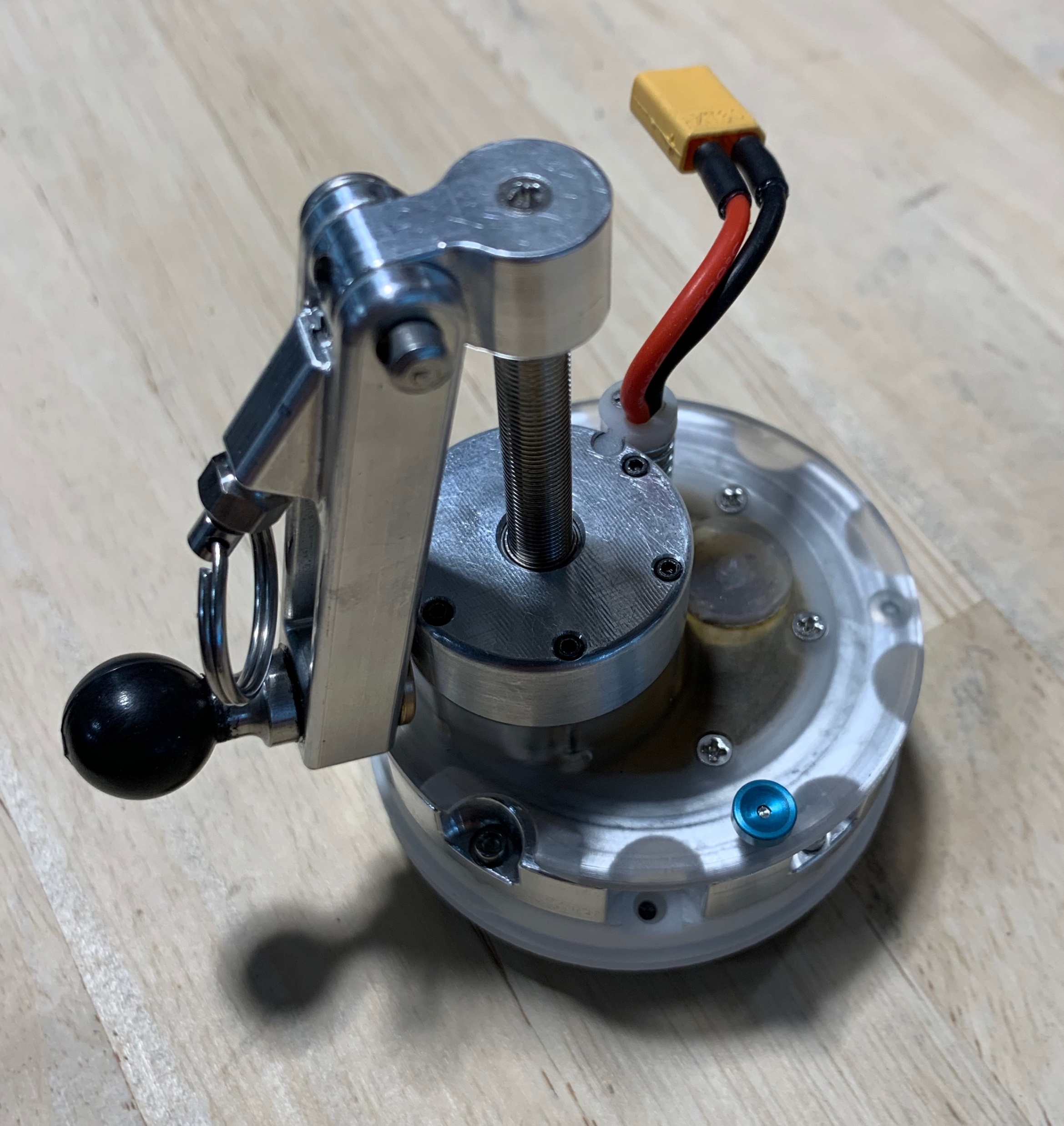

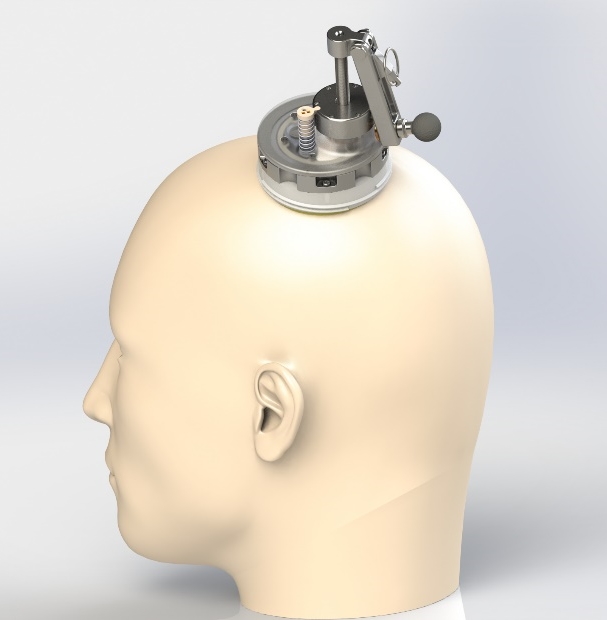

**A**

**B**

**C**

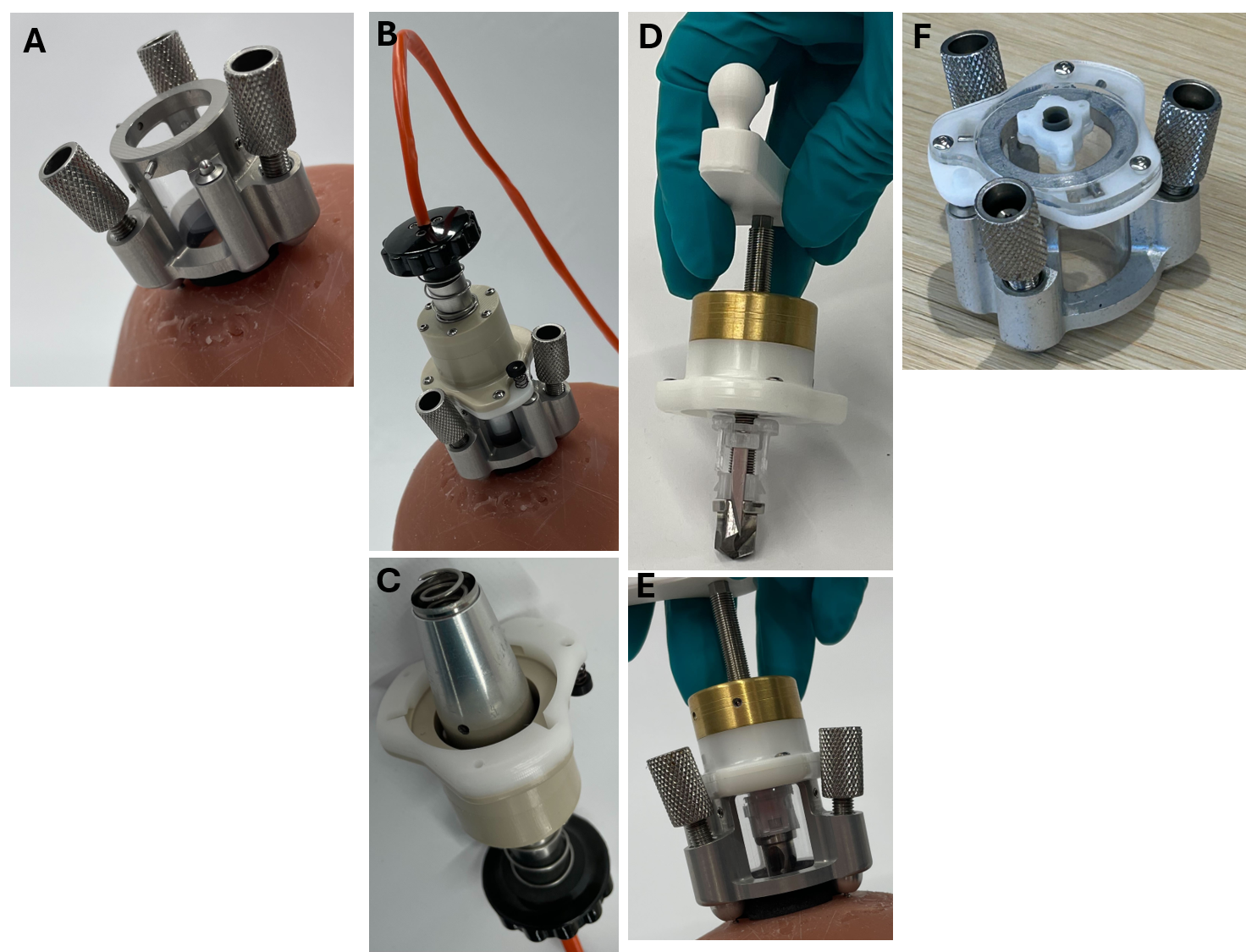

Fig. S13: The design of the IHTD adapted for testing in sheep. The design was constrained by the greater skull curvature in sheep to include a single access port and removable lids for each of the required functions. **A.** The sheep IHTD base affixed to a surrogate via 3 bone screws. **B.** The electrocautery device attached via the orange cable to an external battery and control circuit with feedback temperature control (not shown). In the future, the power and temperature control could be incorporated into the device. **C.** The coiled sharpened hook at the end of the punch cautery to capture the plug of scalp tissue. **D.** The hand crank drill. **E.** The drill deployed through the base of the device. **F.** The intracranial access port.

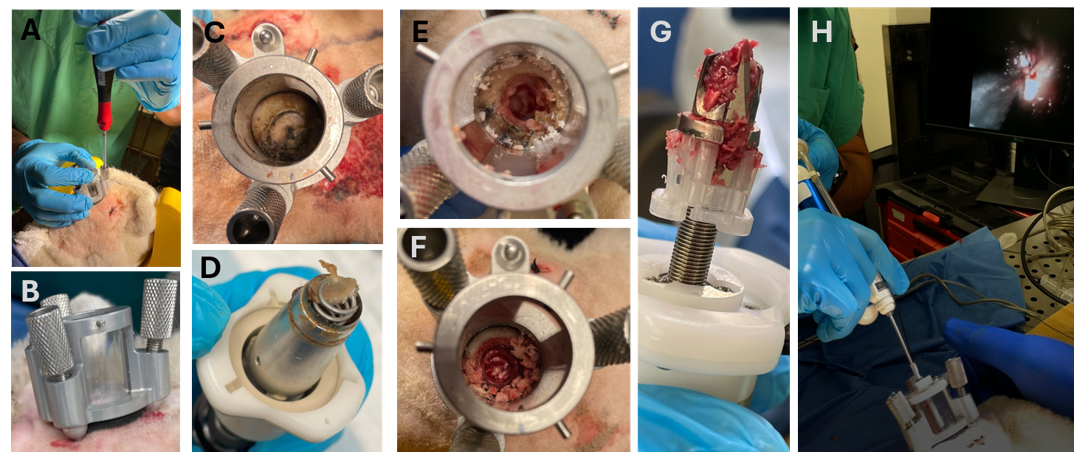

Fig. S14: Use of the Intracranial Hemorrhage Treatment Device in Vivo: **A.** The device was affixed to the previously shaved, cleaned, and sterile-prepped region of the head using 3 bone screws installed using a hand-driven screwdriver. **B.** The device attached tightly, compressing a foam rubber gasket to make a tight seal against the scalp to reduce the risk of infection. **C.** The electrocautery ring burned through the scalp at 232 C to reveal the underlying outer table of the skull. **D.** Rotation of the coiled sharpened screw inside of the electrocautery ring hooked the plug of freed scalp tissue, allowing it to be removed cleanly from the field. **E.** The drill cut through the outer table and inner table of the skull exposing the dura. **F.** The dural cutting blade exposed the subdural hematoma. **G.** The drill and dural cutting blade were removed, bringing bone fragments out of the field. **H.** The subdural spaces was accessed using the Nico Myriad morcellator, with light and camera allowing visualization of the subdural hematoma for morcellation.

##### Randomized Controlled Trial of the IHTD in Sheep

The design of the randomized controlled trial of the IHTD in sheep was prespecified, including the sample size, statistical power, primary outcome measure, and secondary analyses. The prespecified sample size was 12 qualifying sheep, 6 randomized to treatment and 6 randomized to sham. We started with a sample size of 18 sheep because of the natural variability in the model and prespecified that we would exclude approximately 1/3 of the sheep with non-subdural intracranial blood (e.g. intraparenchymal, intraventricular: **Fig. S5-S6**). The prespecified primary outcome measure was reduction in ICP. The trial had statistical power of 0.8 to detect a reduction in ICP with Cohen’s d effect size of 1.5 or greater using a 1-tailed student t-test at p<0.05, and similar power for a 1-sided Mann-Whitney U test in case of non-normally distributed data. Sheep were randomized after the creation of the subdural hematoma and initial ICP measurements to reduce bias. Randomization of 6 sheep per group was accomplished using sealed slips of paper placed in an opaque envelope. Strictly speaking, the last sheep was not randomized because only 1 slip of paper was remaining in the envelope.

Treatment included removing the device from its sterile packaging, affixing it to the sheep head, cauterizing the scalp, drilling the burr hole, cutting the dura, and morcellating the subdural clotted blood. One sheep died after randomization from a respiratory disorder that was determined by the attending veterinarian and veterinary pathologist at autopsy to have been clearly unrelated to any neurological condition or intracranial procedure. Accordingly, this sheep was disqualified. Five other sheep were disqualified before randomization due to intracranial bleeding in non-subdural locations.

Sham involved removing a 1 cm region of scalp by electrocautery and making 3 additional scalp holes to mimic screw holes. There was no burr hole created in the sheep assigned to sham. All sheep had identical appearing sutures and bandages and had the same duration of anesthesia.

Comparison between groups in neurological status post anesthesia and at 24 hours were prespecified secondary analyses. Neurological assessments were performed by 3 board certified veterinarians using a 32-point scale (**Fig. S15**). The 32-point scale was preferred by the USUHS Veterinary Medicine team over the 24-point scale used during model development. The evaluations were performed at 3 time-points: 1) Baseline prior to anesthesia to ensure that all sheep were neurologically normal at baseline. 2) Post-anesthesia, typically 3-4 hours after propofol was stopped, when in the judgement of the veterinarians, the neurological status would reflect the brain injury and not the effects of anesthesia. As noted, veterinarians were blinded to the randomization assignment, the intracranial pressure measurements, and other details of the procedure. 3) Twenty-four hours +/- 30 minutes after the end of the subdural hematoma creation. Longer neurological observations at USUHS were not possible for logistical reasons. The same veterinarian performed all 3 assessments on each sheep. Veterinarians coordinated their assessment strategies in order to homogenize their scoring schema.

Analysis of change in subdural blood volume on CT scans between groups, i.e. from pre-treatment to post-treatment vs. pre-treatment to post-sham was a prespecified secondary analysis. These analyses did not prove to be technically feasible.

Histopathological analyses were prespecified as exploratory analyses. The results of these analyses will be reported separately.

##### Acquisition of CT Scans

Four CT scans were obtained on each sheep.

1) At baseline, to ensure that there were no anatomical abnormalities prior to creating the SDH.
2) After creation of the SDH to localize the blood and exclude sheep for whom the blood was not primarily subdural. This occurred in 6 of the 18 sheep. See **Fig. S5-6** for examples.
3) After treatment or sham to measure the amount of blood removed, vs. natural history of the evolution of blood signal in the brain.
4) Post-mortem.

CT scans were performed in sheep under propofol anesthesia using a NeuroLogica BodyTom portable full body 32-slice scanner leased from Medco Blue. The X-ray tube voltage was 120 kV. A horizonal and a sagittal scout were performed to define the field of view and restrict the CT field of view to brain only. The slice thickness was 1.25 mm. CT scans typically involved 60-70 slices. The reconstruction matrix size was 512x512 pixels. The in-plane resolution was 17-line pairs per cm. The scans required under 3 minutes. Scans were stored on DVDs in DICOM format.

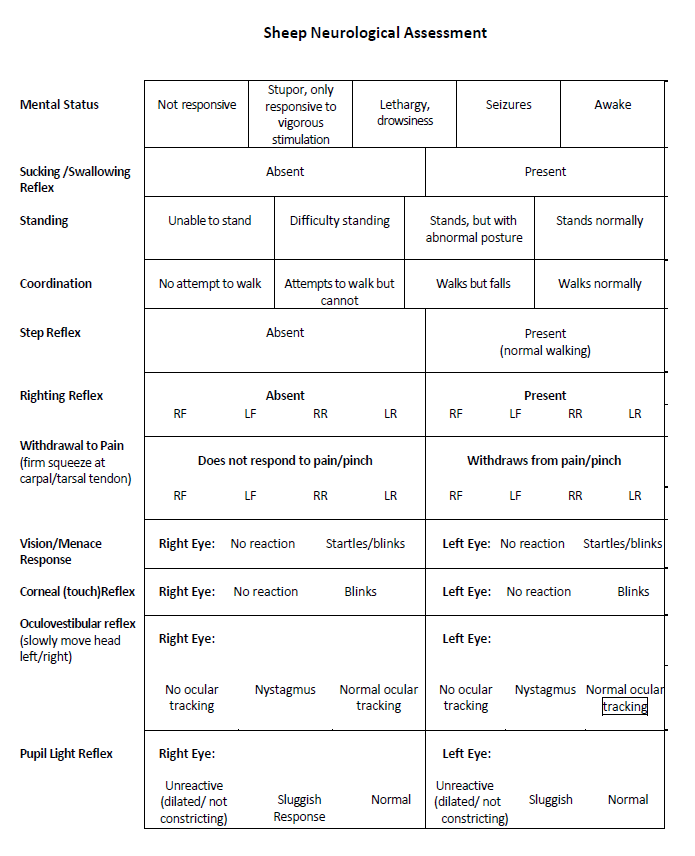

Fig. S15. Sheep neurological assessment using the 32-point scale.

##### Exemplar ICP Tracings During Treatment

An exemplar ICP tracing during the course of treatment with the IHTD is shown in **Fig. S16**. There were stepwise increases in ICP with each infusion of 1 mL of blood, followed by relaxation of ICP. The steady state was reached approximately 20 min after completion of the SDH. There was some variability in ICP during positioning of the sheep for the CT scan. The drill procedure starting around 40 min caused minor, transient increases in ICP. The ICP did not drop until the dura was cut and then dropped further during and after morcellation.

Positioning for CT

Fig. S16 Exemplar ICP tracing from a sheep treated with the IHTD

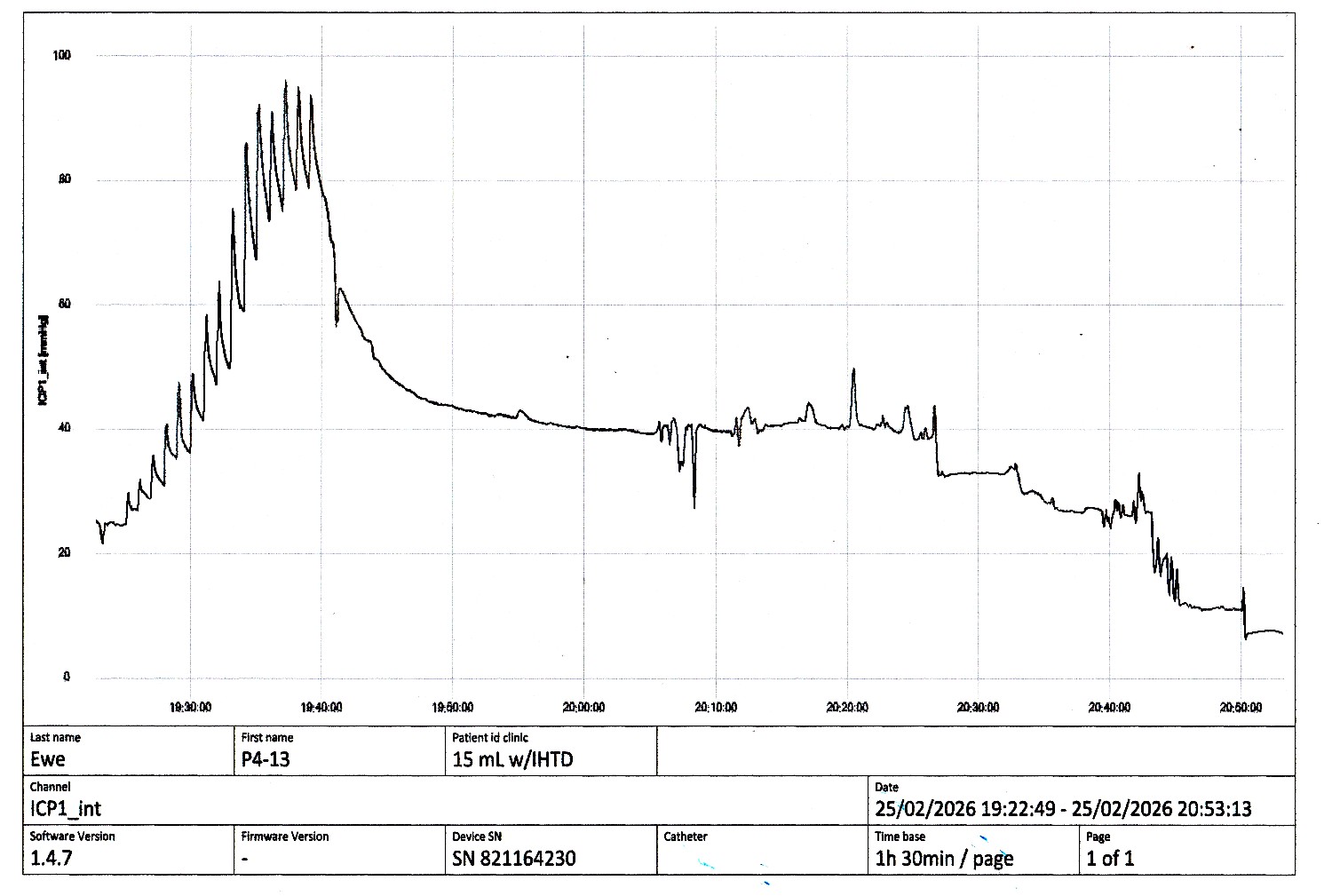

Dural puncture

Morcellation of clot

Drill

##### Correlations Between ICP and Neurological Outcomes

There was a strong and statistically significant correlation between final ICP (approximately 60 minutes after SDH creation) and post-anesthesia neurological outcome (Spearman r= -0.77, p=0.0046), (**Fig. S17**). This result was expected because the final measured ICP was likely to reflect the sustained ICP during the recovery from anesthesia and while they were being assessed neurologically. There were no other correlations with ICPs at other times including peak ICP after subdural creation (Spearman r= -0.23, p=0.46), +20 minutes (Spearman r= -0.47, p=0.12), or +40 minutes (Spearman r= -0.25, p=0.43). There was no correlation with change in ICP following treatment or sham (Spearman r= -0.51, p=0.096).

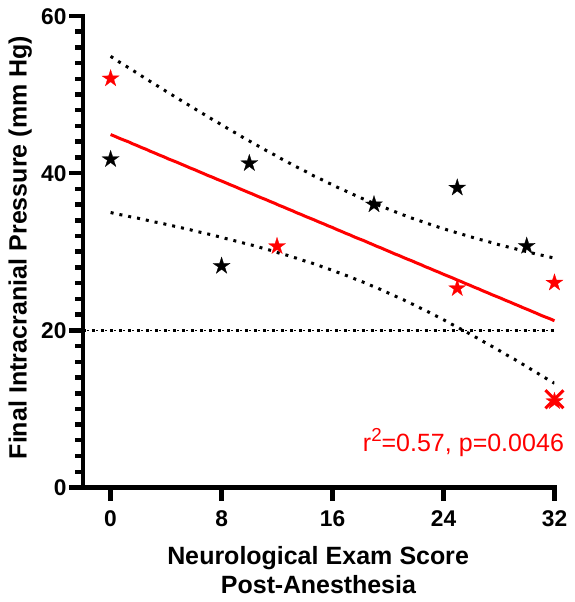

Fig. S17 Correlation between final ICP (approximately 60 minutes after SDH creation) and post-anesthesia neurological status, assessed by blinded veterinarians. Treated sheep are indicated by red stars, and untreated sheep indicated by black stars. The sheep that died before they could be assessed were given a score of 0.

### SUPPLEMENTARY DISCUSSION

As part of these investigations, we have developed a sheep model of severe life-threatening SDH. The relevant characteristics of the model include intracranial pressure elevation, Cushing-like hemodynamic responses, approximately 50% mortality, and neurological impairments in the survivors. Like clinical SDH, there was substantial case-to-case variability at the physiological, neurological, and anatomical levels. This variability represents both a strength and a limitation of the model. Our results indicate a highly non-linear compliance in the sheep brain during infusion of blood into the subdural space. This non-linear compliance and compensatory reserve have been well-described in the human clinical literature (*50*) (*51*). The sheep model may allow for detailed future mechanistic investigations of the pathophysiology of these non-linearities.

There were several limitations of the sheep model. First, only female sheep were used. Male sheep can be more difficult to handle for the veterinary team and can have more irregular skull anatomy. Second, all our studies were performed in young adult sheep, which is appropriate for a military setting but less relevant for civilian settings in which many patients with SDH are elderly. Early surgery is associated with improved outcomes after SDH in elderly patients (*27*), and a potential future direction could involve extending the model to older sheep with age-related comorbidities. Third, we studied only acute SDH. Chronic SDH is also an important clinical problem, but further refinement of methods will be required to produce a valid model. Fourth, the model involved isolated SDH without the contusions, traumatic axonal injury, and other pathologies that can accompany severe TBI in humans. It is not clear whether partially removing subdural clot would have similar clinical benefits in humans with more complex injuries. Fifth, the model involved only a single insult, whereas many patients with TBI sustain multiple injuries including SDH. For example, up to 20% of victims of intimate partner violence sustain SDH, typically in the context of repeated TBI, non-fatal strangulation, and other injuries (*52*). Finally, all of these studies were performed with sheep under propofol anesthesia. For ethical reasons, it is not possible to induce life-threatening subdural hematoma in unanesthetized animals. However, alternative forms of anesthesia should be tested in future experiments. Propofol is convenient and safe, but has been associated with neuroprotective effects (*53*). Thus, the actual injury severity may have been underestimated by our analyses.

The development of the sheep model provides several opportunities for future directions. The sheep model may provide an opportunity for non-neurological surgeons and other military medical providers to practice definitive surgical treatments such as craniotomy and decompressive craniectomy in a controlled fashion prior to deployment. Many military physicians who are not neurosurgeons do not have sufficient familiarity and comfort with these potentially life-saving procedures, and there will always be a relatively small number of deployed neurosurgeons. Importantly, the current pre-deployment training consists of the Emergency War Surgery Course and the ASSETT+ course. Live pigs are used in the Emergency War Surgery Course to replicate abdominal trauma and control life-threatening hemorrhages. The craniotomy portion of the training consists mainly of practice on models and cadavers. The sheep model would be the first living model available for training to diagnose and treat life-threatening intracranial hemorrhage of which we are aware. In civilian settings, general surgeons can treat severe TBI quickly, but mortality may be higher (*54*). Training with a large animal model has potential for benefit by improving speed and confidence. By analogy to acute ischemic stroke, faster time to definitive treatment may result in better outcomes. In a recent survey, time from injury to emergency surgery for TBI averaged 13 hours (*55*), which is far slower than the optimal time to treat brain ischemia from stroke within 1 hour (*56*). Additional future potential uses for the model include assessments of new neuro ICU care strategies, investigations of relevant aspects of cerebral physiology including autoregulation under extreme elevations of intracranial pressure, preclinical trials of neuroprotective medications, and development of novel surgical procedures, and many others.

There are several limitations of the findings involving the detection and treatment devices. First, the devices have been tested only in a single large animal model of subdural hematoma, and not in humans. While it is likely that the detection and treatment devices will be just as useful for epidural hematoma as for subdural hematoma, testing in epidural hematoma should be explicitly performed. Second, the heads of the sheep used were closely shaved and had light colored skin; the accuracy of the detection device in the presence darker colored skin has not been explicitly assessed. Shaving the head in humans to allow optimal accuracy for the detection device is reasonable when there is life-threatening severe traumatic brain injury. It is possible that the currently marketed detection devices such as the Infrascanner 2000 which do not require shaving may be more appropriate for triage use in less severe injuries. Third, for logistical reasons, we were not able to continue neurological evaluations during the randomized controlled trial for more than 24 hours. Likewise, we have not assessed other neurological outcomes relevant to traumatic brain injury such as memory, social behavior, headache-related behavior, and sleep. Fourth, only a single non-neurosurgeon used the treatment device during the randomized controlled trial; multiple medical personnel with the skills and education of the intended users will need to be trained to use the treatment device. Fifth, we have not explicitly tested the longer-term stability of the treatment device when anchored to the skull, nor the effectiveness of the seal designed to reduce the risk of infection. Thus, these should be considered the first in a series of experiments required to address unmet needs related to life-threatening traumatic brain injury when CT scanners and neurosurgeons are not available.

While promising, considerable additional work will be required before these devices are ready for human use. First, the laboratory prototype devices will need to be miniaturized and ruggedized for field use. All the components are available in miniaturized form, and none are especially delicate or temperature sensitive. Second, the new detection device will need to be tested in humans with traumatic brain injury, using CT scan results are the ground truth. With miniaturization and validation in humans, it is likely that the new detection device could be approved quickly by the FDA under the 510(k) program. Third, the version of the treatment device intended for human use will need to be constructed and tested in human head surrogates and cadavers. Fourth, the treatment device will need to be tested under Good Laboratory Practices standards by an independent research organization for replication and external validation. Fifth, criteria for appropriate use of the treatment device will need to be developed for FDA approval for the treatment device, likely under the Class III Premarket Approval pathway since to our knowledge a predicate device does not exist. Finally, larger scale manufacturing, distribution, storage, training, maintenance, and replacement conditions for the devices will need to be established for them to be incorporated into routine practice.

If successfully transitioned to human use, there are many potential scenarios where these devices could be employed, either individually or together.

1) Both devices: In large scale combat operations involving peer or near-peer adversaries or special operations, forward military medical providers could be trained to use the detection and treatment devices as part of tactical combat casualty care and prolonged field care. It is likely that these devices would be used in role 2 facilities such as battalion aide stations, after the most immediately life-threatening injuries such as massive bleeding, airway compromise, respiratory failure, and cardiac arrest have been treated by Role 1 first line providers. In other truly remote, austere, or hazardous conditions where no medical evacuation is possible, the detection and treatment devices could be used together by on-site medical personnel to assess and when appropriate provide initial treatment to patients with life-threatening traumatic brain injury. Such patients are typically in a deep coma and will have prolonged amnesia if they survive, thus the risk of pain or distress caused by performing a burr hole is likely to be outweighed by the risk of death or profound neurological disability from the injuries themselves. As noted above, the largest domain for potential use is likely to be in the developing world where severe traumatic brain injury is most common, while CT scanners and neurosurgical facilities are most scarce. Even when facilities are available, the cost of a CT scan is out of reach for many people in the developing world. The long-term burden of traumatic brain injuries is comparable to that of infectious diseases in many parts of the world. However, much less philanthropic support has been provided to address traumatic brain injuries, perhaps in large part because technological solutions have not been developed to address the unmet needs. The long-term costs to society in terms of lost lives and lost productivity likely will vastly outweigh the shorter-term costs of the devices and training.
2) Detection device only: In prehospital environments where some limited medical evacuation is possible, the detection device could be used to help make an assessment about whether medical evacuation is warranted. For example, it may be worthwhile to send a helicopter team into a dangerous environment when there is a clearly treatable and time-critical life-threatening subdural or epidural hematoma, but not worthwhile under other circumstances. In mass casualty situations involving multiple patients with severe traumatic brain injury, the detection device could be used to determine which patients could most benefit from direct transportation to a neurosurgical facility. The receiving neurosurgical team could also be alerted in advance to reduce time to definitive treatment.
3) Treatment device only: In trauma centers with CT scanners but without neurosurgical facilities, emergency medicine providers and other non-neurosurgeons could use the treatment device to initially mitigate the elevated intracranial pressure caused by subdural and epidural hematomas prior to transfer for definitive neurosurgical care.
